# The global geometry of optimal neural representations

**DOI:** 10.64898/2026.11.17.767450

**Authors:** Gengshuo John Tian, Brent Doiron

## Abstract

Efficient coding hypothesizes that neural activity optimally represents information under resource constraints. The global structures of such optimal population codes are difficult to study theoretically, and previous works have usually resorted to either focusing on local properties like fine discrimination or placing restrictions on the parametric forms of neural tuning. We address the general problem by analytically deriving the neural representation that minimizes the all-pairs average classification error of a one-dimensional circular variable under various forms of neural response noise. Our method is based on formulating the space of neural response distributions as a statistical manifold whose geometry can be simplified through a generalized version of variance-stabilizing transformations. The theory fully characterizes the complete family of optimal population codes and allows us to make two novel predictions. First, an optimal code should allow the existence of oppositely peaked bimodal tuning curves. Second, there should be a correlation between spike count variability and tuning sharpness. We validate both of these predictions in the mouse head direction system.

## Introduction

Subjective experience is markedly different from physical measurements, with the degree of perceptual similarity between stimuli defining a geometry over an internal perceptual space [1]. From a neuroscience perspective, the perceptual space is biologically instantiated by neural activity. Characterizing and understanding the geometry of these neural representations is thus a central goal of systems and computational neuroscience, as it opens a window into the organization of the brain’s internal model of the world [2, 3].

A longstanding challenge in the study of perceptual spaces is that even though local discriminability is relatively well characterized, the global organization of the space remains much less well understood. In psychophysics, distinct methods have been developed to measure sensory discrimination thresholds [4] and suprathreshold perceptual differences [5], but the relationship between these two types of measurement remains an important open question [6]. In neuroscience, rapid progress in experimental techniques has enabled the collection of large datasets of population neural activity, but extracting the behaviorally relevant structures from data has become an increasingly challenging task [7]. Decoding analysis on neural data has achieved notable successes in explaining perceptual decision task behavior near threshold [8], but the global organization of the neural code in relation to perceptual experience remains less well understood [9]. These challenges highlight the need for forming mathematical theories that establish principled relationships between local perceptual sensitivity, global perceptual structure, and their underlying neural representations [10].

A promising approach is to construct normative theories of discriminability, which aim to understand neural representations as being optimized for computational goals expressed as objective functions [11–13]. The canonical objective quantifying the local discriminability of continuous variables is Fisher information (FI), and there has been extensive work on finding the population codes that maximize it under models of neural variability [14– 17]. On the other hand, while measures such as mutual information (MI) from the efficient coding framework [11, 18–21] are global objectives, they are notoriously difficult to study analytically [9]. Consequently, previous works have often resorted to making restrictive assumptions on the shape of tuning curves such as bell-shaped bumps to make the problem tractable [14, 15, 20, 22–24]. However, non-canonical tuning curve shapes outside of the presumed constraints are frequently observed in experiments [25–28], and different assumptions may lead to seemingly contradictory predictions about the optimal representation [21, 29].

In this paper, we present a novel theoretical framework that takes steps to meet this challenge. We consider the representation of a variable by a noisy population of neurons optimized for discriminability, with optimality measured by the average discrimination error between all pairs of stimuli. We also consider the function space of all smooth population tuning curves with no restrictions on the functional form of the neural code. In particular, we focus on one-dimensional periodic variables, which frequently appear in neuroscience experiments as drifting grating orientation [30], coherent visual motion direction [31], direction of arm movements [32], and head direction [33]. We analytically derive the complete family of optimal solutions under various neural variability models.

Our theory produces two predictions for the optimal code, one concerning the range of tuning curve shapes and the other relating neurons’ variability to their tuning sharpness. Both predictions are then tested in a head-direction cell dataset from freely moving mice [34]. We also extend our result to multi-dimensional variables including spheres and tori. The central insight underlying our result is that there is a correspondence between the structure of neural variability and the geometry of the ambient manifold where neural representations reside. In sum, our results lay the foundation for a new theoretical framework on the global geometry of optimal neural representations.

## Results

### A global cost function for discriminability

We consider in this study the representation of a circular variable *θ* ∈ [0, 2*π*], which can be sensory, motor, or cognitive. Neural activity exhibits considerable fluctuations across trials with the same experimental conditions [35–38]. As such, we describe the representation of *θ* by a population of *N* neurons as a map *f* from *θ* to the *distribution* of neural response *f* (*θ*) = *p*(*x* | *θ*), where *x* ∈ ℝ^*N*^ is the vector of spike counts. For stimulus values *θ* and *θ*^′^, the overlap between the distributions *p*(*x*|*θ*) and *p*(*x* | *θ*^′^) leads to potential confusion between the two. Therefore, a representation *f* that optimizes for discriminability should be organized such that the amount of overlap is minimized.

FI is commonly used as a measure of discriminability for neural codes. It is related to the classification error between *θ* and *θ*^′^ = *θ* + Δ*θ* for small Δ*θ* [14], and can be thought of as a differential version of the sensitivity index *d*^′^ (“d-prime”) widely used in perceptual studies [39]. However, FI fails to account for coarse discrimination performance, i.e., when Δ*θ* is not small [22, 40, 41]. Consider a toy example with *N* = 2 neurons with tuning curves *r*_*i*_(*θ*) = E[*x*_*i*_ | *θ*] (*i* = 1, 2) and whose variability is i.i.d. Gaussian (Fig. 1a, point clouds) [9]. We shall use *r*(*θ*) to denote the collection of tuning curves in the population, also called the population tuning curve. We compare the discriminability of response geometries *r*(*θ*) that go from a round circle to a narrow ellipse, all parameterized by arc length (Fig. 1a, red closed curves). The three codes have the same FI (by design), but MI, which is a global measure of coding, decreases as the eccentricity of the ellipse increases (Fig. 1a, inserts; Methods). In line with this, the classification error between *θ* and *θ* + Δ*θ* is virtually identical for all three geometries for small Δ*θ* (Fig. 1b, blue), but the chance of *global* errors for large Δ*θ* increases dramatically for a narrow ellipse compared to a round circle (Fig. 1b, green). This is because *r*(*θ*) and *r*(*θ* + Δ*θ*) can be embedded much closer together in the ambient neural space for the elliptical code compared to the circular code (Fig. 1a, green), leading to a larger overlap between the corresponding point clouds. This structure is not captured by FI, which is only sensitive to *local* errors (Fig. 1a, blue). Since confusion over larger Δ*θ*’s has conceivably graver behavioral consequences, these global errors are sometimes referred to as catastrophic errors [41, 42]. It is therefore critical for the neural code to avoid them as much as possible.

**Fig. 1.**
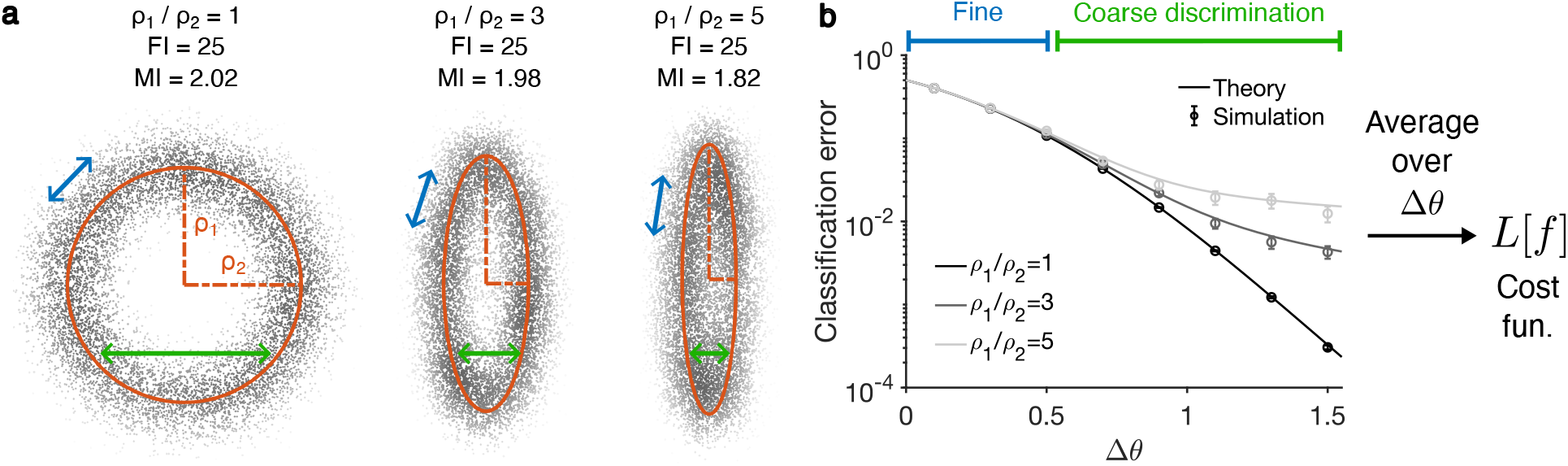
**a**, Three example neural codes of *θ* ∈ [0, 2*π*] by 2 neurons with isotropic Gaussian noise. From left to right the population tuning curves are ellipses with aspect ratios 1, 3, and 5 respectively and whose lengths are held constant, leading to constant Fisher information (FI) but decreasing mutual information (MI). Illustration inspired by [9]. **b**, Neurometric curves of the three codes in **a**, with theoretical predictions by Eq. (2) matching simulation results. The three codes have similar performances in the fine discrimination regime where the two *θ*’s needing classification are close to each other, but have drastically different performances in the coarse discrimination regime. Error bars are standard errors of the mean (SEM). The total error (or equivalently the average error) is the area under the curve, which is the cost function *L*[*f*] we use in this paper.

Motivated by the above example, we consider a global cost function for the code *f* defined as:

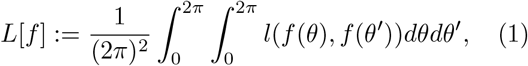

where *l*(*f* (*θ*), *f* (*θ*^′^))) is the lowest achievable classification error between the two distributions *f* (*θ*) = *p*(*x*|*θ*) and *f* (*θ*^′^) = *p*(*x*|*θ*^′^). *L*[*f*] can equivalently be defined as the area under the neurometric curve (Fig. 1b), which incorporates both fine and coarse discrimination. To emphasize the global structure of the neural code over its local structure, we fix the total FI of *f* which sets the overall available coding resource or model complexity [9, 21, 29, 43– 45]. For simplicity, we assume for now that *θ* is uniformly distributed, which according to efficient coding principles requires the magnitude of FI to be constant over *θ* [21, 29].

### The optimal population code for binary classification

Our central goal is to derive the optimal neural code, namely the *f* that minimizes *L*[*f*] under the appropriate constraints. When neural variability is i.i.d. Gaussian with constant variance, as in our toy example (Fig. 1), the classification error *l* between stimuli *θ* and *θ*^′^ is solely determined by the distance between the mean responses *r*(*θ*) and *r*(*θ*^′^). In that case, we can directly invoke an existing result [46] to conclude that the circle is the minimizer of *L*[*f*] (Fig. 1a, left), which is intuitive given the symmetry of the problem.

However, the problem is more complicated for general variability structures because the classification error is no longer a function of the mean response difference alone. Consider, for instance, a neuron with Poisson variability. Due to trial-to-trial variability growing proportionally with the firing rate, neural response pairs with the same *difference* in their means now have larger variability overlaps when the *absolute values* of their means are large. Consequently, a population tuning curve in the shape of a circle no longer minimizes *L*[*f*], because stimulus pairs with the same Δ*θ* still evoke the same difference in mean response (which is the chord length of the circle), but the classifier error now changes along the circle (Fig. 2a), breaking the symmetry required for optimality. Intuitively, then, we need an alternative notion of distance that is statistical in nature and describes the amount of overlap between neural response distributions *f* (*θ*). Such a distance would then be inhomogeneous with respect to the mean response parameterization, a phenomenon that is classically studied in differential geometry.

**Fig. 2.**
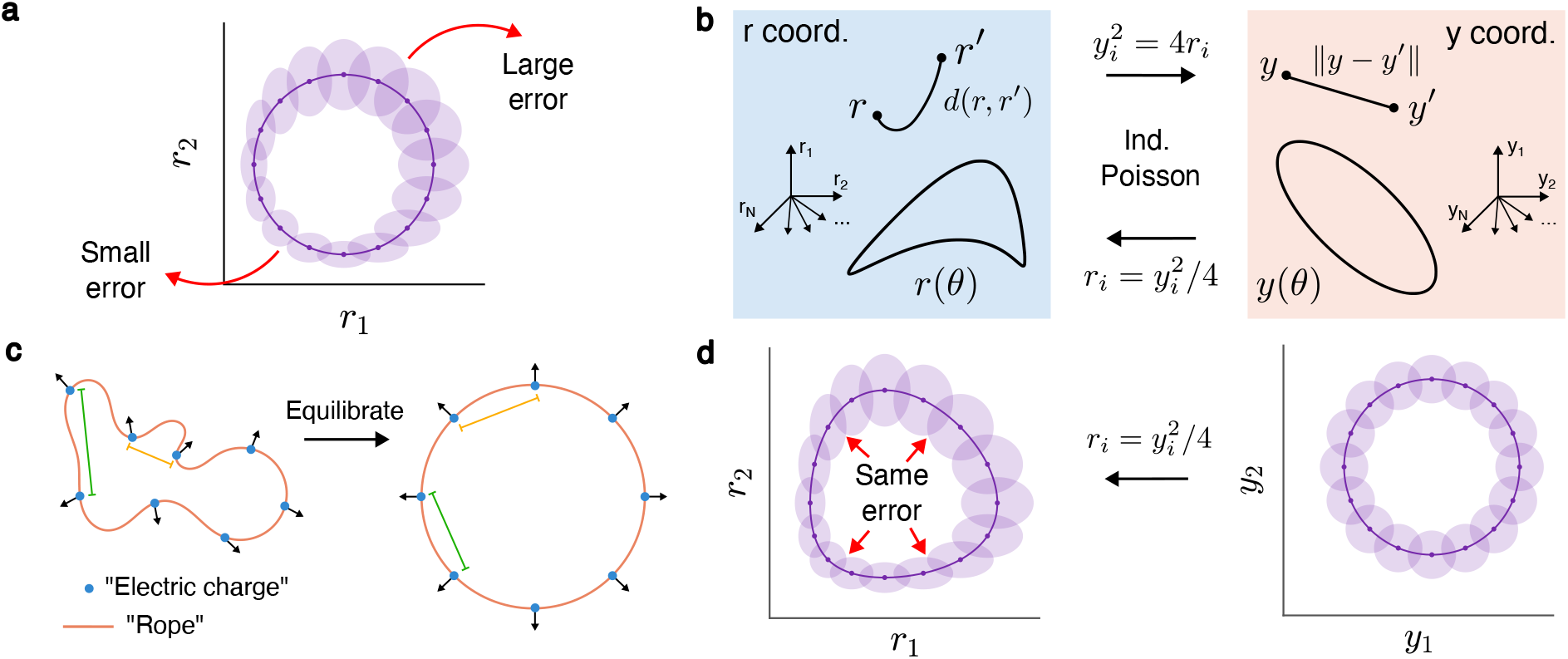
**a**, A circular population tuning curve in firing rate *r* coordinates is not optimal under Poisson variability. In particular, there is an uneven distribution of classification errors across *θ*’s, shown in the plot as overlaps between ellipses that delineate the noise covariances around evenly spaced *θ*’s by covering 80% of the probability. **b**, The mean-noise relationship defines a Riemannian metric in the space of neural response distributions parameterized by the firing rate *r* (left, blue). The induced geodesic distance *d*(*r, r*^*′*^) is difficult to calculate. Transforming to an alternative *y* coordinates (right, red) makes the metric Euclidean. The optimization problem is easier in *y* coordinates, and the solution *y*(*θ*) can be transformed to the population tuning curve *r*(*θ*) by reversing the transformation. The *y*-*r* transformation is determined by the noise model. The independent Poisson case is shown as an example. **c**, Physical intuition of why the circle optimizes *L*[*y*]. For *h*(*d*) = 1*/d*, the cost function can be thought of as the total energy potential from Coulomb-like repelling interactions between electric charges (points) uniformly fixed on the curve *y*(*θ*) that decrease with distance, with its fixed length being analogous to the rigidity of a rope. (There is a technical point concerning integrability that can be addressed by regularizing the functional into the Möbius energy [51, 52].) As the collection of charges reaches equilibrium and the potential is minimized, the forces balance each other out through the symmetry of the circle. Although some distances between pairs of charges decrease while others increase, the overall average interaction energy is minimized. **d**, The solution curve that minimizes the cost function is a circle in *y* coordinates (right). It is no longer a circle when transformed back to the firing rate *r* coordinates (left), but statistical properties such as the symmetry of classification errors is retained.

We now formalize this idea using independent Poisson neurons as an example. Observe that the spike count distribution *p*(*x*) is fully determined by the firing rate vector *r* = E[*x*] due to the independent Poisson assumption. This implies that the space of neural response distributions is a *N* - dimensional manifold parameterized by *r*, which allows us to replace the map of conditional densities *f* (*θ*) by the simpler population tuning curve *r*(*θ*) without any loss of generality. The statistical distance discussed earlier can then be formalized as the Fisher-Rao distance (which we denote by *d*) from information geometry (Supplementary Information A.1), making the statistical manifold a Riemannian one with metric *g*^*r*^ [47, 48]. The relation between this distance and the classification error *l* has been made precise by previous work [14]:

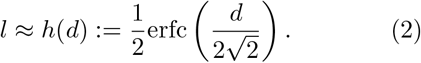

Additionally, under the geometry we defined, the total FI is now simply the length of the curve *r*(*θ*) [29]. Then the feasible set for the cost function *L*[*r*] becomes all smooth closed curves on the manifold with the same length. We shall not impose any additional assumptions on the functional form of *r*, making the minimization of *L*[*r*] a variational problem, in contrast to most previous works that restrict the shape of the tuning curves to be a function described by a few parameters, most commonly bell shaped curves [15, 20, 22–24].

For i.i.d. Gaussian noise with constant variance, the metric *g*^*r*^ is simply Euclidean, but in general *g*^*r*^ takes a nonlinear form due to the dependence of variability on the mean. Under independent Poisson variability, the metric is 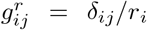 (see Eq. (S6)). For such a *g*^*r*^ the minimization of the cost function *L*[*r*] is intractable (blue box in Fig. 2b). However, we can transform the problem by using an alternative coordinate system 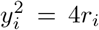, which makes the Fisher-Rao distance *d* explicitly Euclidean in the new *y* coordinates (red box in Fig. 2b). This is closely related to the variance-stabilizing square root transform commonly used to analyze spiking data [49, 50] (see Discussions). We can then invoke the result [46] which states that the minimizer of *L*[*y*] is the circle (see theorems 1 and 2 in Supplementary Information A.2). This completely characterizes the family of solutions to the discriminability problem: all circles of the same radius in *y* coordinates have the same optimal performance, and all optimal codes take this circle form.

To gain some physical intuition for why the circle minimizes *L*[*y*], consider the alternative integrand *h*(*d*) = 1*/d* [45], which is also convex and decreasing. Then *L*[*y*] becomes the Coulomb potential of a collection of electric charges tethered on a rigid closed rope, where each pair of points on the rope repel one another with a force that falls off with distance (Fig. 2c, left). While the electric force pushes points away from each other, the tension along the rope prevents unbounded separation and keeps the total length constant. The average *h*(*d*) is minimized when these forces balance each other, and this occurs when the rope is shaped as a circle (Fig. 2c, right). Note that this symmetric structure hinges on the ambient metric being Euclidean, highlighting the criticality of the transformation into *y* coordinates.

To obtain the optimal population tuning curve *r*(*θ*) from the circle *y*(*θ*), we need only reverse the coordinate transformation given earlier (Fig. 2d). In the case of independent Poisson neurons, this reverse transformation is an element-wise square, and we thus call the optimal code the “square-of-circle” (SoC) code. The resulting population tuning curve appears warped, but the curving is matched to the ambient geometry such that the statistical properties of the neural code are symmetric. Indeed, for fixed Δ*θ* the classifier error is independent of the location on the curve (compare Fig. 2d to Fig. 2a). Changing the noise model alters the geometry of the neural space and leads to different coordinate transformations and ultimately different optimal tuning curves. We shall explore the consequences of different neural variability structures in later sections.

There is a small caveat due to potential sign ambiguities from the squaring transformation for some circle configurations, but its effect is shown to be vanishing in large population size *N* (Supplementary Information A.3). This ambiguity is not present when *y*(*θ*) is nonnegative, and we shall use this strictly optimal code unless otherwise stated.

Interestingly, the theorems in [46] allow us to draw even stronger conclusions in two respects. First, the minimizer of the functional *L*[*y*] is the circle for any function *h* that is convex and decreasing, not just the specific one given in Eq. (2). This implies that even if the real neural population is not optimizing over the average pairwise classification error, as long as the cost function can be written as the average of some convex decreasing function of pairwise Fisher-Rao distances on the curve, the solution to the problem is the same one.

Put in another way, the same neural representation is optimal for a wide range of processing needs whose cost functions take this general form. This is encouraging, as the downstream task presumably differs across receiving brain regions and changes with context, and our result suggests a converging optimal representation for a large class of tasks [53]. Second, the circle (in *y* coordinates) does not just have the smallest overall classification error, but also has the best performance for *each* Δ*θ* after averaging over (*θ, θ* + Δ*θ*) pairs. That is, its neurometric curve is uniformly optimal regardless of how fine or coarse the discrimination task is.

Note that requiring the curve *r*(*θ*) to be smooth and closed amounts to restricting its *topology* (assuming *N >* 3), which can be detected in data using topological data analysis (TDA) [54]. Our theory builds on top of this assumption and instead focuses on the *geometry* of the code, with the circle being a geometric statement expressing the underlying symmetries of the problem as well as the neural space. The topology of the encoded variable is critical. If *θ* ∈ [0, 2*π*] is not periodic and instead an open-ended interval, then the topology of *r*(*θ*) is that of an arc, and the optimal code becomes a straight line segment in *y* coordinates, which leads to a quadratic population tuning curve for independent Poisson noise.

### Heterogeneous tuning curves

We next evaluate the SoC code by comparing it to the classical von Mises (VM) model with unimodal and symmetric tuning curves that uniformly tile the *θ* space [41, 45, 55–57] (Supplementary Information A.4). VM tuning curves have identical shapes (Fig. 3a, top left) which lead to the same selectivity across the population (Fig. 3a, middle left; see Methods for the definition of selectivity). In contrast, the SoC code exhibits heterogeneous tuning shapes with a wide range of selectivities (Fig. 3a, top and middle right), which is more aligned with observations from neural data [26, 27]. Note that this heterogeneity does not play a functional role as in [24, 26, 42, 58], but is instead an expression of the degrees of freedom in the solution set [59, 60]. Underlying this apparent heterogeneity lies a simple two-dimensional circle at the population level in *y* coordinates, in contrast to the higher-dimensional and somewhat more complex curve from the seemingly more regular, yet suboptimal, VM model (Fig. 3a, bottom).

**Fig. 3.**
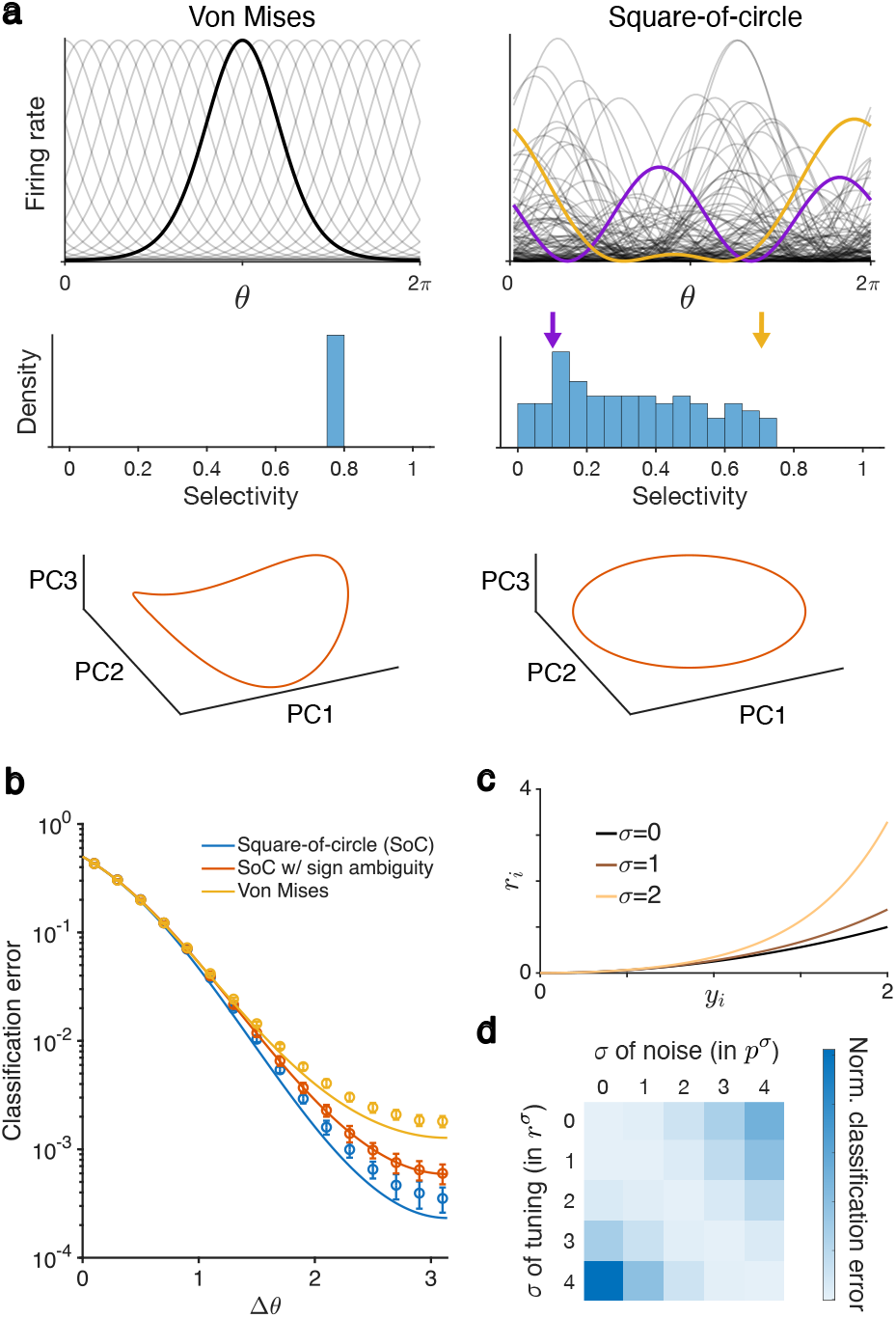
**a**, Top: Example tuning curves from the classical von Mises model and the square-of-circle (SoC) code. Middle: Distribution of selectivity for the two codes. Bottom: The top 3 principal components (PCs) of the two codes’ population tuning curves in *y* coordinates. The VM model occupies more than 2 linear dimensions, in contrast to the 2-dimensional circle in the SoC code. **b**, Theoretical predictions and simulation results for the neurometric curves of the von Mises code (Eq. (S18)), the nonnegative SoC code (Eq. (S11)), and the SoC code with sign ambiguity (Eq. (S13)). Error bars are standard deviations. **c**, The transformation between *r* and *y* coordinates under Gamma-Poisson noise indexed by *σ* (Eq. (S21)). **d**, The performance of the neural code when the *σ* parameter in the noise structure (*p*^*σ*^) and the *σ* parameter in the tuning curves (*r*^*σ*^) vary. The classification error is only minimized when the two are matched according to the theory presented in Supplementary Information A.5.

As predicted, simulations show the SoC code significantly outperforming the VM model in coarse discrimination while having similar performance in fine discrimination (Fig. 3b). Here, the parameters are set such that the two codes have the same total FI and average maximum firing rate (Methods). The SoC code’s superiority is maintained even if moderate sign ambiguity is present (Fig. 3b, compare red and yellow curves). Further, the tuning curves in the SoC code are all less selective than the VM model (Fig. 3a), consistent with the observation in previous work that a neuron’s selectivity is a poor predictor of its importance in coding [61].

We emphasize that the transformation 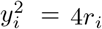 used in the SoC code directly arises from the assumed independent Poisson noise structure. A different mean-noise relationship would lead to a different metric *g*^*r*^, which in turn would require a different transformation to the Euclidean *y* coordinates. To demonstrate this point, we consider the Gamma-modulated Poisson model [38] where the firing rate of each neuron *i* is multiplicatively modulated by a Gamma variable *G*_*i*_ with mean 1 and variance *σ*^2^ that is independent across neurons and trials, i.e.,

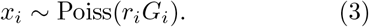

This gives a family of mean-noise relations parameterized by *σ*, which we denote with *p*^*σ*^. Different values of *σ* lead to different coordinate transformations to Euclidean coordinates (Fig. 3c) [62]:

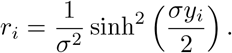

This in turn defines different optimal codes, again parameterized by *σ*, which we denote with *r*^*σ*^ (Supplementary Information A.5). We can pair a noise model *p*^*σ*^ with a population tuning curve 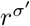 with various *σ*^′^ values to get different combined neural codes. Our theory predicts that only when the tuning curve and the variability structure match — that is, only when *σ*^′^ = *σ* — is the representation optimal. This is indeed verified in simulations (Fig. 3d; Methods).

In sum, we have presented a framework that finds the population tuning curve *r*(*θ*) which minimizes the average classification error *L*[*r*] among all smooth curves, and in this sense produces an optimal neural code. The resulting neural representation not only outperforms classical models in coarse discrimination but also generates tuning curves that exhibit heterogeneity closer to what is observed in neuronal data. We next examine two specific predictions of our theory on several features of population response.

### Signatures of the optimal code observed in head-direction cells

Head-direction (HD) cells, found in several areas of the mammalian brain, have been extensively studied [33]. HD is a one-dimensional circular variable that is densely sampled by freely moving animals. For these reasons, the HD system is distinctly suitable for testing our theory. In the following, we verify several predictions of our model in a recent neurophysiology dataset recorded from the postsubiculum (PoSub) of mice freely exploring an open arena (Fig. 4a) [34].

**Fig. 4.**
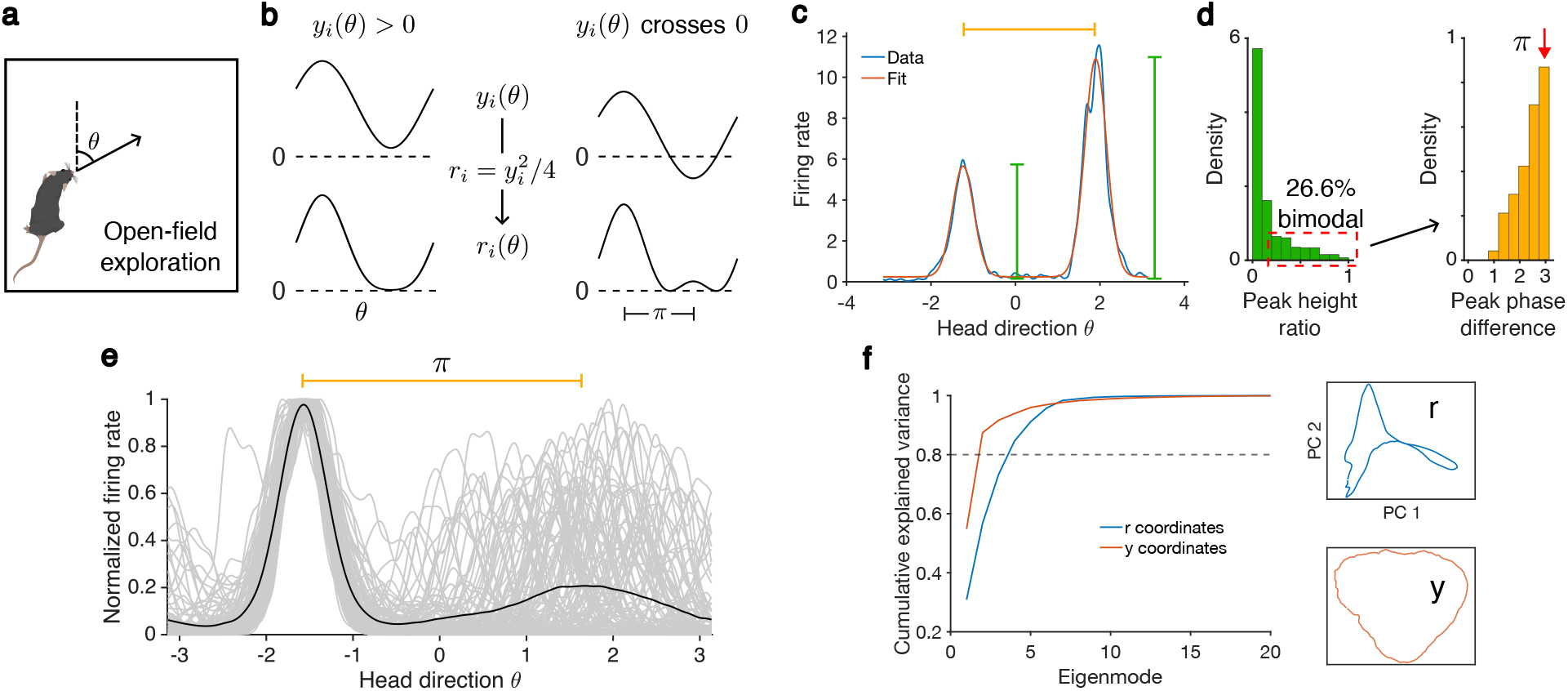
**a**, The dataset in [34] consists of recordings in mouse postsubiculum during free exploration of an open arena. **b**, an SoC tuning curve can be either unimodal or bimodal depending on whether it changes sign in *y* coordinates. For bimodal tuning curves, the two peaks must have a phase difference of *π*. **c**, The tuning curve from an example neuron and the fit to a sum-of-von Mises model, which captures the shape of the two modes. **d**, Left: The distribution of the ratios between the heights of the fitted modes demonstrates that approximately 1/4 of neurons are bimodal (with a cutoff value of 0.2). Right: The distribution of the phase differences between the two peaks of bimodally tuned neurons concentrates around *π*. **e**, Plotted in gray are the bimodal tuning curves as identified in **d** shifted and normalized such that their larger peaks are located at −*π/*2 with magnitude 1. The black curve is their average, which shows a clear second peak at *π/*2. **f**, Left: The (cumulative) scree plot from performing PCA on the population tuning curve in the original firing rate coordinates (*r*) and the transformed Euclidean coordinates (*y*). It has a high linear dimension in *r* coordinates (requiring at least 4 dimensions to explain 80% of the variance) but a low linear dimension in *y* coordinates (requiring only 2 dimensions to explain 80% of the variance). Right: The population tuning curve in the original *r* coordinates and the transformed *y* coordinates visualized in the first 2 PCs.

The first prediction concerns the qualitative shape of the HD cell tuning curves. Let us revisit the structure of tuning curves from an SoC code. A circle in *y* coordinates corresponds to (vertically and horizontally shifted) cosine functions *y*_*i*_(*θ*) for each neuron. After squaring, *r*_*i*_(*θ*) can be either unimodal or bimodal depending on whether *y*_*i*_(*θ*) changes sign (Fig. 4b). For a bimodal tuning curve, the two peaks must have a phase difference of *π* because they come from the maximum and minimum of the cosine in *y*_*i*_(*θ*). While real neurons likely do not have strictly independent Poisson variability, this qualitative prediction still holds for a large family of noise models including the Gamma-Poisson model (Eq. (S21)). Strikingly, bimodality is present in approximately a fourth of the tuning curves from the HD dataset (Fig. 4c, d; Methods), even though there is a priori no obvious computational reason to expect neurons to be tuned to opposite HD. Further, when a HD tuning curve is bimodal the separation between peaks is concentrated near *π* (Fig. 4c-e).

At the population level, our theory predicts that the SoC code *r*(*θ*) should have 4 linear dimensions (Methods), and transforming it into *y*(*θ*) should recover a 2-dimensional object that resembles a circle. This was verified in the data using principal component analysis (PCA), as *r*(*θ*) requires at least 4 dimensions to explain 80% of the variance while *y*(*θ*) virtually occupies only 2 dimensions (Fig. 4f, left). Additionally, the geometry of the first 2 PCs of *r*(*θ*) is not very interpretable, whereas the PCs of *y*(*θ*) form a curve that is close to a circle (Fig. 4f, right), which is a natural representation of HD.

### Neurons with more response variability have sharper tuning curves in an optimal code

A core feature of our theory is the correspondence between the noise model and the optimal population tuning curve (as illustrated in Fig. 3d). While the variability structure of recorded neurons arises from the underlying circuit dynamics and presumably stays constant through the experimental time frame, we can nevertheless test this aspect of the theory by examining how tuning curves vary with the mean-noise relationship across different neurons.

Consider again the Gamma-Poisson model (Eq. (3)), but now allowing the variance 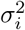 of the modulation variable *G*_*i*_ to be neuron-dependent (in contrast to Fig. 3d, where all neurons have the same *σ* parameter). Then the optimal tuning curve for each neuron *i* should depend on its *σ*_*i*_ (Eq. (S21)). In the Gamma-Poisson model [38], neurons with a larger *σ*_*i*_ show more super-Poisson spiking activity (Fig. 5a; Eq. (S20)). In the corresponding optimal tuning curve formula (Eqs. (S10) and (S21)) a larger *σ*_*i*_ implies a sharper peak (Fig. 5b). Crucially, these two aspects of the neural response can be independently fitted from data, the former from the variance-noise relation and the latter from the tuning curve shape (Methods). We denote the results of these two fitting processes with *σ*_var_ and *σ*_tun_ respectively. From a statistical point of view, *σ*_var_ relates the first- and second-order statistics of spike counts with no regard for the encoded variable *θ*, whereas *σ*_tun_ only concerns how the first-order statistic changes with *θ*, so they are not expected to be related to each other a priori. However, our theory predicts that a correlation should exist in an optimal code. Satisfyingly, *σ*_var_ and *σ*_tun_ are indeed correlated (Fig. 5c), and the result is consistent across all 17 mice in the dataset (Fig. 5d).

**Fig. 5.**
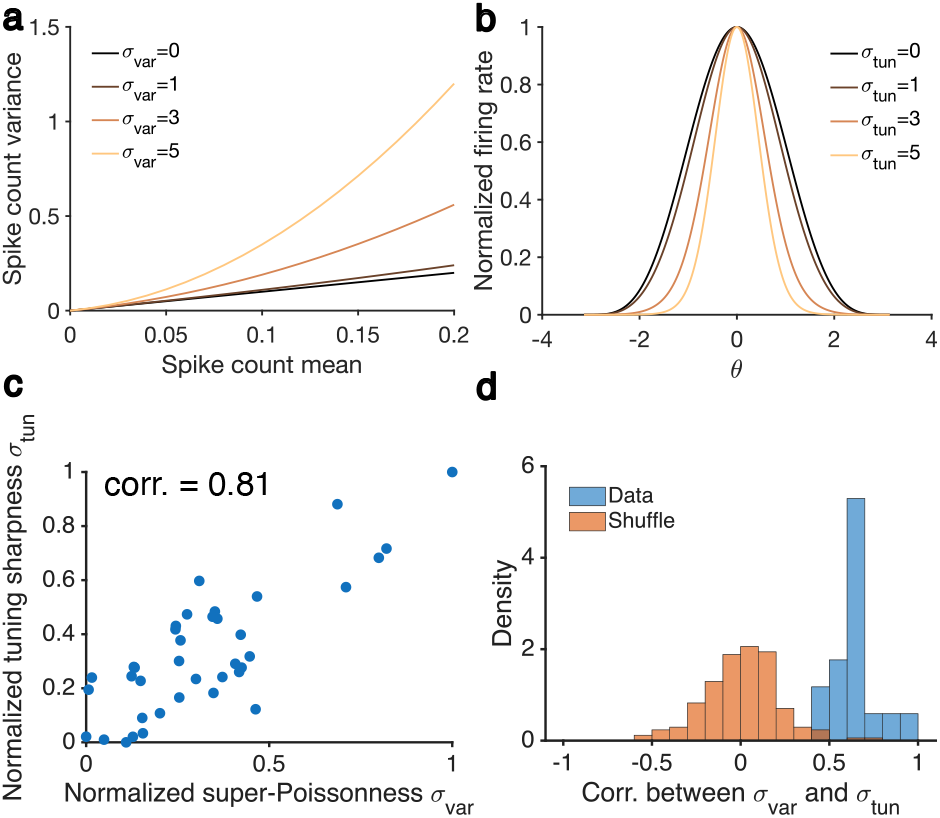
**a**, The variance-mean relationship (Eq. (S20)) as the *σ* parameter in the Gamma-Poisson model changes. **b**, The tuning curve shape (the transformation of a cosine *y*(*θ*) according to Eq. (S21)) as the *σ* parameter in the Gamma-Poisson model changes. The notations *σ*_var_ and *σ*_tun_ are introduced to highlight the fact that these parameters can be estimated from data independently from the variability structure and the tuning curve of the neuron, respectively, even though they are predicted to be equal in our theory. **c**, Data from an example session showing that *σ*_var_ and *σ*_tun_ are correlated. Each point on the scatter plot is data from one neuron. *N* = 85 neurons in total. See Methods for a discussion on the necessity of normalization. **d**, Correlations between the two *σ*’s across 17 mice compared to shuffled data.

### Recurrent connectivity determines the optimal code through noise correlation structure

To apply our framework we require the metric *g*^*r*^ on the response distribution manifold to allow a transformation to an explicitly Euclidean space (the *y* coordinates) where we can employ the theorem in [46]. This is trivially the case when neurons are independent, as the statistical manifold would then be the product of *N* manifolds [48]. It is however less clear if this is the case when there are noise correlations in the neural variability model.

To explore this question, we consider a recurrent circuit model with connectivity matrix *W* receiving external noise *ξ* whose covariance matrix is *C*_0_. When the network can be linearized around a fixed point, the covariance matrix of neural activity has been classically derived to be (Supplementary Information A.6) [63, 64]

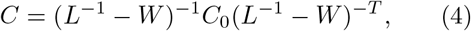

where *L* is the diagonal matrix of neuronal gains. If we assume that neurons have a threshold-quadratic transfer function, then we have the relation *L* ∝ *R*^1*/*2^ where *R* := diag(*r*) [65], which makes the covariance *C* = *C*_*W*_ (*r*) a function of the firing rate *r*. If we additionally assume that the spike count vector follows a multivariate Gaussian distribution *x* ∼ N(*r, C*), then there is again a bijection between the mean response *r* and the full distribution *p*_*r*_(*x*).

If we use linear FI, which measures the discriminability of linear decoders [16, 66], the resulting Riemannian manifold is again Euclidean, and the transformation between *r* and *y* coordinates is expressed by a system of quadratic equations that can be solved numerically (Supplementary Information A.6).

The above reasoning states that the recurrent connectivity of a network determines the optimal representation through the noise correlation structure. As an illustration, we examine random weight matrices with i.i.d. Gaussian entries and with *C*_0_ = *I*. Different realizations of *W* from this distribution lead to different mean-noise relationships *C*_*W*_ (*r*), which in turn produce different transformations that take the circle *y*(*θ*) to the optimal population code *r*(*θ*) (Fig. 6a). In other words, the same circle *y*(*θ*) transforms into different (optimal) tuning curves for different connectivity matrices (Fig. 6b). We can again investigate the importance of matching variability (defined through *W*) and tuning with simulations. As predicted by our theory, the classification error is only minimized when the tuning curves are matched with the specific realization of *W* that set the variability structure (Fig. 6c). This is true even though all connectivity matrices are generated by the same underlying distribution.

**Fig. 6.**
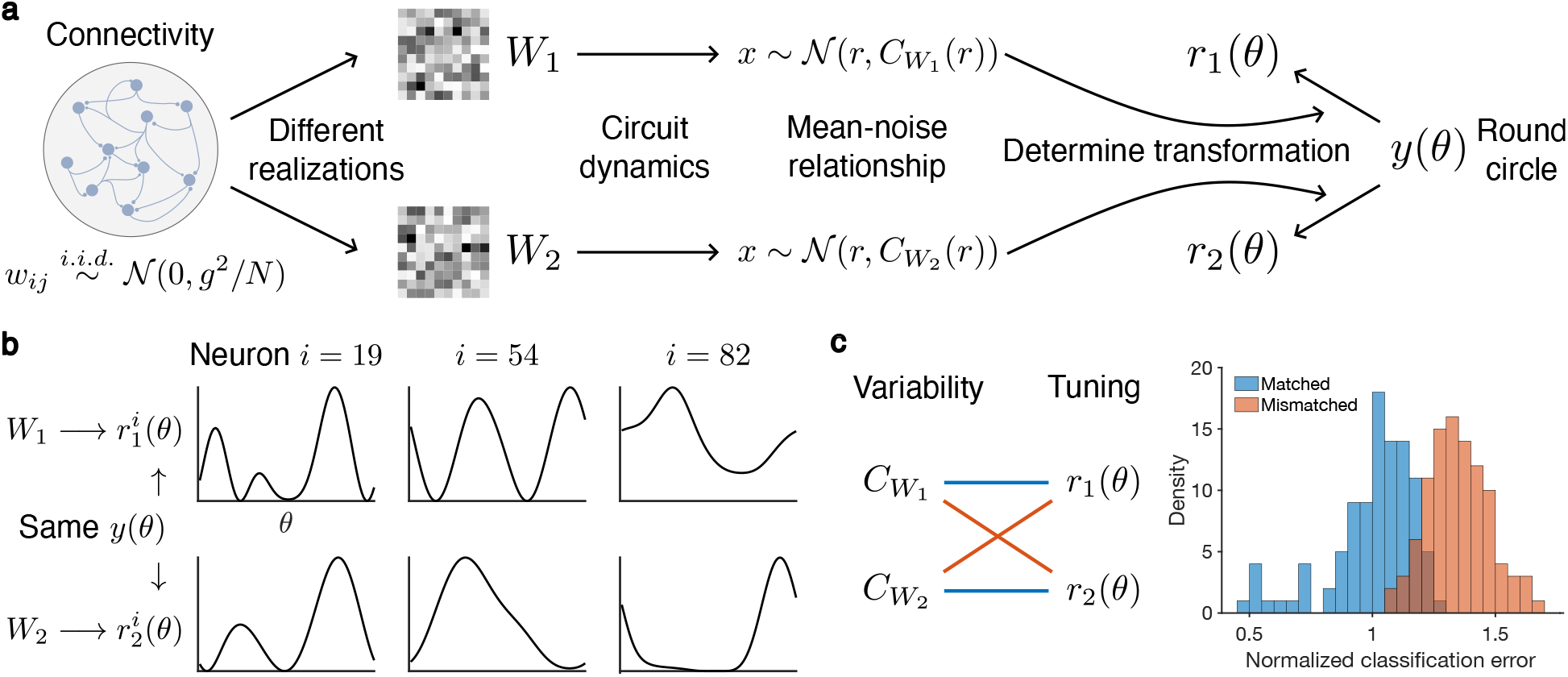
**a**, Different realizations of the recurrent connectivity lead to different mean-noise relationships, which in turn produces different optimal tuning curves. **b**, Optimal tuning curves of 3 example neurons under two realizations of recurrent wiring. **c**, Simulation result showing that performance is significantly worse when the tuning curves are not matched to the connectivity matrix that generates the response variability. The histograms are across 100 realizations. Classification errors are normalized such that the mean error of the matched case is 1.

Here, we only simulated the i.i.d. Gaussian weight matrix as a first demonstration of the value of our framework in understanding the coding properties of recurrent circuits. Different structures in the connectivity *W*, the neuronal gain *L*, and the external noise *C*_0_ can all potentially generate novel features in the optimal code. Crucially, obtaining the optimal tuning curve can be carried out without much difficulty by numerically solving Eq. (S24), making it a promising direction to explore in future work.

### Global gain modulation produces hyperbolic geometry

Another way to introduce noise correlations into the system is with shared top-down modulations. Specifically, one may consider the multiplicative gains *G*_*i*_ in Eq. (3) to have some correlations with each other [38]. In the extreme case of perfect correlation, we have a random variable *G* that is globally applied to the population (i.e., *G*_*i*_ = *G* for all *i*). Under this variability model, the FI metric is no longer Euclidean, but instead hyperbolic with a constant negative curvature. We prove that the optimal population tuning curve here is the element-wise square of an ellipse under mild assumptions, which is qualitatively indistinguishable from the SoC code (Supplementary Information A.7). Nevertheless, intermediate levels of structured correlation in the gain modulation are expected to produce nontrivial geometry that is not Euclidean, offering an alternative approach to modeling noise correlation.

### Additional energy constraints introduce a trade-off between discriminability and robustness

For neural codes with the same discriminability performance, the one with smaller firing rates may be more desirable since it consumes less energy [67]. This can be included in the optimization problem by constraining the population firing rate Σ*r*_*i*_ to be constant for all *θ* (Fig. 7a, top, gray region). This constraint can also be viewed as the result of normalization [68, 69]. Under the independent Poisson noise model, this amounts to constraining the curve *y*(*θ*) to lie on a sphere centered at the origin (Fig. 7a, bottom). When the total FI constraint (i.e., the length of *y*(*θ*)) is too large, the optimization problem is no longer analytically solvable (Supplementary Information A.8).

**Fig. 7.**
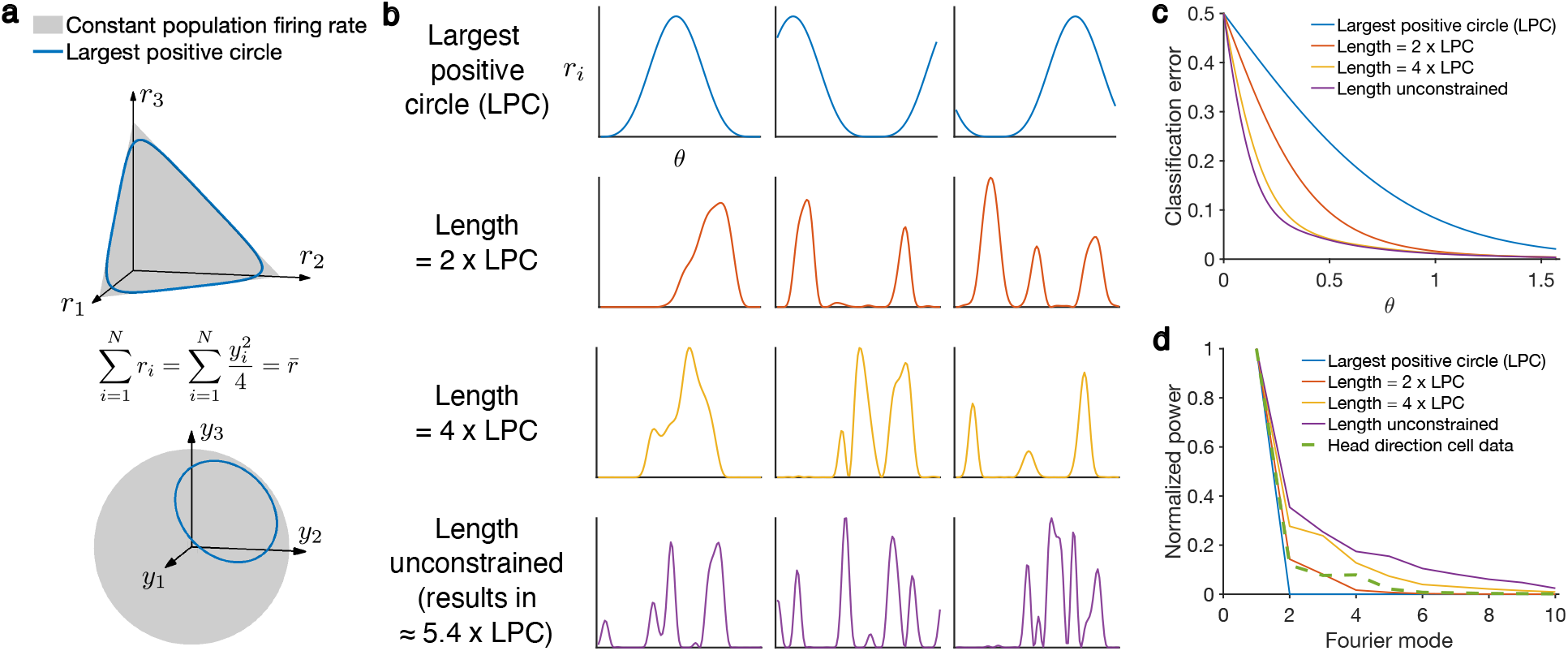
**a**, Schematic showing the manifold on which the population tuning curve is constrained in *r* (top, which is a simplex) and *y* (bottom, which is a sphere) coordinates. The largest positive circle (LPC) is the largest circle on the manifold where all points have nonnegative elements, which is the reference point for our simulations as the length constraint is gradually relaxed. **b**, Example tuning curves as the length of the curve is allowed to be longer and longer in each consecutive row until it is unconstrained in the bottom row (except the monotonicity requirement Eq. (S26)), which results in a length of around 5.4 that of LPC after optimization. **c**, Neurometric curves of the optimized tuning curves in **b. d**, Fourier spectra of the optimized tuning curves (in *y* coordinates) in **b** as well as those in the HD dataset, averaged over the population.

We thus turn to numerical optimization (Methods). The resulting tuning curves become less and less smooth as *y*(*θ*) is allowed to become increasingly longer (Fig. 7b). While a larger total FI does lead to better discriminability (Fig. 7c), the oscillatory nature of the code reduces its robustness since the shape of the tuning curves is highly fine-tuned and any small perturbation to the system or the input may drastically affect its performance. A code with larger curvature would also make it harder for downstream decoders to generalize, as it would take more data to capture the microscopic details of the representation, and the complexity of the decoder would be required to match that of the code. Intuitively, the compactness of the sphere on which the curve *y*(*θ*) is constrained forces the curve to be more convoluted (i.e. less smooth) as its length increases, which affects its robustness and generalization properties. This is consistent with the total FI being a measure of model complexity [43, 44].

One way to evaluate the smoothness of the code is to look at the power of the larger Fourier modes of the tuning curves. In *y* coordinates, circles have no power at the second Fourier mode and above. As the total FI increases, the larger modes of the optimal code gain more and more power (Fig. 7d). Interestingly, the HD cell data falls on an intermediate state, suggesting that both the total FI constraint and the energy constraint should be taken into account when understanding the neural representation, and that there is a trade-off between discriminability and robustness [70].

A recent work also explored the coding implications of a compact ambient manifold [42], but in the context of random compression. They concluded that there is an optimal level of smoothness that balances local and global errors. In contrast, in our case both local and global errors decrease as the code becomes less smooth due to the active optimization process (Fig. 7c), and the trade-off is instead between discriminability and robustness. Another work performed a similar optimization [45], but with a different energy and a weaker constraint (Supplementary Information A.8). They observed similar improvements in performance and reduction in smoothness as the curve length is increased. The bounded nature of the available neural space has also been invoked to explain the curved representation for decision variables in monkey lateral intraparietal cortex (LIP) [71].

### Non-uniform prior distribution of the encoded variable

The optimal code should depend on the prior distribution of the encoded variable *θ*, as is evident in the formula for MI. Up to now, we have assumed a uniform prior in our derivations. When the prior *p*(*θ*) is not uniform, which encodes for example the more frequent occurrence of horizontal and vertical orientations in natural scenes [72], the cost function Eq. (1) should be weighted by the pairwise frequency *p*(*θ*)*p*(*θ*^′^). Additionally, efficient coding principles suggest that the square root of FI, which in our formulation is exactly the speed along the curve, should be proportional to the prior [20, 21, 29]. In this case, a rescaling of *θ* recovers the original problem and results in a code with the same circular shape but whose speed is reparameterized (Supplementary Information A.9).

More generally, we may consider the frequency of *θ* pairs presented in the hypothetical experiment to have a joint distribution *p*(*θ, θ*^′^). If the probability of each pair depends only on their difference, then the theorem in [46] states that the optimizer is again the circle in *y* coordinates. This case can be thought of as placing a weight on the relative importance of fine and coarse discrimination. Interestingly, the optimal code does not depend on this weighting if the marginals *p*(*θ*) and *p*(*θ*^′^) remain equal.

### Representing multi-dimensional variables

A natural extension of our result is to consider the representation of multi-dimensional variables. We prove that the optimal representation for hyperspheres and hypercubic flat hypertori is the inclusion into *y* space when the embedding does not allow local stretches (Supplementary Information A.10). However, it is unclear whether this assumption is neuroscientifically justified, and there is considerable nuance in properly defining the optimization problem. More generally, the global minimizers for other multi-dimensional *θ* manifolds are unfortunately unknown [73]. However, qualitative characterizations of the solution set are still valuable and potentially feasible [74], and optimization algorithms are also available for numerical explorations [52].

We have focused on HD cell data up to now. There are many other settings in neuroscience where circular variables are represented, most notably orientation selectivity in the primary visual cortex (V1) [75]. However, it is not obvious whether our theory is applicable to this scenario, because V1 collectively represents the entire visual scene, which is quintessentially high-dimensional, and even single neurons respond to many features other than orientation [26, 76, 77]. This calls into question whether we should frame a V1 population as optimizing solely for encoding orientation. In contrast, it has been argued that the main function of PoSub is to calculate and represent HD signal [78].

We nevertheless tested our theory on a V1 dataset where monkeys passively view drifting gratings with *θ* being the drift direction [30]. Since there is a natural confounding factor of neurons encoding the *direction* versus the *orientation* of the drifting gratings, it is difficult to test our predictions on the bimodal shape of tuning curves (Fig. 4a-e) and we focus instead on the relation between variability and tuning sharpness (Fig. 5). Interestingly, the predicted correlation between *σ*_var_ and *σ*_tun_ still holds (Fig. S2). This suggests that V1’s representation of *θ* as a one-dimensional submanifold of the full visual space may be close to optimal.

If we maintain that V1 codes for the much higher-dimensional visual space, then this raises the question of when an optimal code for a multi-dimensional variable is also optimal for a submanifold and what this reveals about these particular directions in the stimulus space. A concrete theory addressing this question will immensely expand the applicability of our model, as a typical neuroscience experiment only explores a small subset of the variables involved in the recorded population’s full range of computations.

## Discussion

In the paper, we proposed the average binary classification error as a global measure of the discriminability of neural representations (Fig. 1). The minimizers of this cost function for a one-dimensional periodic variable *θ* ∈ [0, 2*π*] are obtained by characterizing the space of neural response distributions as a Riemannian manifold whose geometry depends on the mean-noise relationship of the population response (Figs. 2, 3). The theory predicts that an optimal code should allow the inclusion of neurons whose tuning curves are bimodal with peaks separated by *π*, and that increased super-Poisson variability for a neuron should be correlated with an increased sharpness of its tuning curve. Both of these predictions are novel to our knowledge, and are verified in a HD cell dataset [34] (Figs. 4, 5). The developed technique can also be applied when there are noise correlations from recurrent interactions and external inputs to elucidate how these factors affect the optimal code (Fig. 6). Finally, numerical explorations of additional energy constraints reveal a trade-off between discriminability and robustness (Fig. 7).

### Additional considerations are required to explain other features of the representation

A curious consequence of our result is that all circles *y*(*θ*) with the same length have the same mean binary classification error regardless of the location of their centers and which 2-dimensional subspace they occupy (besides the slight caveat of sign ambiguities that was addressed in Supplementary Information A.3). Then there are many features of tuning curves that are not accounted for by the normative theory we presented and need additional constraints to explain [9]. For instance, if the circle is contained in a 2D coordinate subspace, then only 2 neurons participate in this code, which is highly unusual. However, to achieve the same FI value as *N* neurons with *O*(1) firing rates, these 2 neurons would need firing rates of order *O*(*N*), so a constraint on the biophysical properties of individual neurons would lead to a more uniform spread of both activity level and tuning across neurons. Another consideration might be that downstream decoders typically only have access to a subpopulation of neurons. This incentivizes the code to be near optimal in each random subpopulation, which can be achieved by generating each neuron’s tuning curve in a statistically identical manner [59]. The strength of our framework is that it disentangles discriminability from other objectives, isolating their different effects on the representation.

Even with additional constraints, it is conceivable that the reduced solution set may still consist of a continuum of possibilities. It has been proposed that representational drift, where neural tuning changes slowly over a timescale of days or weeks, can be understood as a random walk on the manifold of equivalent population codes [79]. Thus, a characterization of the full set of optimal codes may help uncover the underlying principles behind the observed drift patterns.

### A canonical geometry over the neural response space

We defined a canonical geometry on the manifold of neural response distributions (Fig. 2a). This allows us to reformulate statistical questions about the neural code in geometric terms [9]. In the particular setting of encoding a one-dimensional periodic variable, we expressed the optimality of the code as the geometric shape of circles (in *y* coordinates).

Many theories and data analysis methods implicitly place a flat geometry in firing rate space by either assuming mean-independent noise or disregarding variability altogether [45, 80–84]. For instance, Euclidean distance and linear correlation are often used in representational similarity analysis [81]. One reason for this is that in order to model the global structure of neural representations, a space with a globally defined geometry is required to connect the local structures. Such a definition cannot be extrapolated from limited data samples of the space without exceptionally strong assumptions, and the Euclidean assumption is arguably the reasonable default choice. The same can be said about modeling perceptual spaces as well. Our framework, however, suggests that a principled choice of Riemannian geometry in the neural space can be made from the structure of neural variability.

Our results further suggest that recognizing this geometry yields important insights and predictions. A compact illustration of this point is the fact that a *l*^1^ constraint in the original *r* coordinates (i.e., requiring the sum of firing rates to be bounded) becomes a *l*^2^ constraint in the explicitly Euclidean *y* coordinates (i.e., requiring the code to lie inside a ball, Fig. 7a) for independent Poisson neurons. These different regularizations have distinct predictions on features of neural representation like sparsity [85]. Adapting existing theories and analysis methods to the proposed canonical geometry is thus a promising direction of research.

The mathematical study of this geometry is made accessible in large part by the transformation from the firing rate vector *r* to an alternative coordinate *y* where the FI metric is explicitly Euclidean (Fig. 2b). This procedure gives rise to a space where neural variability is isotropic. Then the symmetries in the neural representation manifest as familiar geometric structures in the Euclidean space (Fig. 2c,d). In particular, many features of the neural code such as discriminability are then invariant to Euclidean isometries (i.e., translations, rotations, and reflections). In the other direction, the representation in the original firing rate space is sensitive to isometries. Recent work has discussed why there are privileged axes in biological and artificial neural networks [86], and our framework provides an additional explanation for this phenomenon.

A geometric understanding of the neural code’s statistical properties can also be used to interpret changes of the representation across states and contexts. For example, multiplicative modulations of tuning curves have often been observed, whether due to contrast [87], locomotion [88], or attentional state [89]. In the SoC code, this amounts to increasing the total FI while preserving the code’s optimality for the given FI values. In contrast, additive shifts to tuning curves do not preserve the population code’s shape and break optimality, coinciding with the conclusion of another model [90].

The core idea of seeking an isometrically noisy space for neural response is not new. A transformation that is virtually identical to Eq. (S21) was employed in [62] as a step in the estimation of the neural trajectory’s curvature. More broadly, variance-stabilizing transformations have long been employed in the analysis of spiking data [9, 49, 50], with the square-root transform for Poisson data being the most common. However, note that these transformations are applied directly to the (trial-averaged) spike count *x* instead of the mean response *r*. The contribution of our framework is that we circumscribed the conditions for such a transformation’s existence, namely the FI metric being Euclidean. This allows us to generalize the transformation method to a large family of variability structures including noise correlations in linearizable recurrent circuits (Eq. (4)).

### Relating neural space to perceptual space

It is a long-standing question in the study of perception how sensory thresholds, which measure local discriminability, relate to suprathreshold perceptual distances [6]. Fechner hypothesized back in the 19th century that perceptual intensity, which is a form of perceptual distance, can be obtained by integrating local sensitivity [91], but recent empirical evidence suggests that the perceptual geometries inferred from threshold and suprathreshold judgments can be inconsistent [92]. It has been proposed that the perceptual color space may be non-Riemannian in nature [93].

Our theory presents a framework for relating local discriminability and global distances in perception. We view neural representation as a submanifold of the full *N* -dimensional statistical manifold of neural response distributions, which for the main case considered in this paper is simply a closed curve in Euclidean space. Then perceptual thresholds measure the local metric (i.e., FI) in the direction neural activity changes in response to stimulus change [16], while perceptual distances correspond to the global distance in the ambient neural space. The latter depends on how the representation is embedded in the manifold of all neural activity patterns, and the same local sensitivity profile can correspond to different suprathreshold properties depending on the embedding.

According to our theory, an assumption of optimality provides a unique embedding at least for circular variables. It then predicts that the perceptual distance between two stimuli separated by Δ*θ* should be proportional to 2 sin(Δ*θ/*2), which is the chord length on a circle. This specific proposal is not new [94], and many other functions can take a similar qualitative form, making it difficult to test rigorously in data [93]. However, our theory provides a general recipe for generating predictions on perceptual distances that is rooted in neural response geometry and efficient coding principles, which has the potential to broadly unify threshold-based and suprathreshold measurements. For example, our theory can be applied to multidimensional variables such as spheres and tori (Supplementary Information A.10), which can be used to generate predictions on perceptual distances in future work.

### The variability-tuning relation as a bridge for understanding neural data

The variability structure of real neural populations is often exceedingly complicated and may change across contexts and brain states. Characterizing it with statistical models has thus been a challenging and ongoing direction of research [95].

On the one hand, our theory suggests that a finer understanding of neural variability is needed to derive the optimal tuning curves more accurately (Fig. 3d and Fig. 6c). Adopting better noise models yields a closer match to data. For example, using the Gamma-Poisson model to fit the tuning curve results in an average *R*^2^ value of 77% compared to 57% when the SoC model (which is based on independent Poisson neurons) is used. Incorporating noise correlations will likely generate additional insights into neural data (Fig. 6).

The derivation of the optimal code requires knowledge of the variability around every firing rate vector *r* ∈ ℝ^*N*^, but we can only probe those around the population tuning curve *r*(*θ*). This issue can potentially be mediated by working within a certain class of models such as Eq. (4), utilizing data from other brain states such as spontaneous activity, and using better statistical methods such as a recent one based on Wishart processes [96]. Our results motivate seeking a better understanding of the full space of neural variability, which requires both new statistical methods and experimental designs that focus on trial-to-trial fluctuations.

On the other hand, our theory can also transfer population tuning curve geometry to knowledge about neural variability. Given an existing noise model (that is perhaps supported by previous datasets) and under the assumption that the neural representation is close to optimal, one can use the correlations implied by our theory (such as those in Fig. 5) to infer the variability structure of the recorded neurons from their tuning curves (e.g., obtaining *σ*_*var*_ from *σ*_*tun*_). Since estimating the mean response requires much less data than characterizing the trial-to-trial fluctuations directly, this approach may allow for the exploration of neural variability structures under richer experimental designs and stricter constraints.

### Mechanistic models of bimodal tuning curves in the HD system

Bimodal tuning curves whose peaks are predominantly separated by a phase difference of *π* have been reported before for HD cells in the medial entorhinal cortex (MEC) [97, 98]. The HD system is classically modeled as a ring attractor [99, 100], as it maintains a stable representation of HD information even in the absence of sensory stimuli and can continuously update it [33]. Correspondingly, there has been work studying double-peaked patterns in ring attractor networks. One work found a large region of parameters where the only stable pattern is double-peaked [101]. Another studied how the number of bumps in the activity pattern affects the computational properties of the network and found that having multiple bumps has negligible effect on noise-driven deviations in the bump position [102]. How this dynamical perspective interacts with our results is an interesting topic for future research.

## Methods

### Simulations

#### Ellipses with isotropic Gaussian noise

In Fig. 1, we illustrated the local nature of FI by simulating three population codes of 2 neurons whose mean responses *r*(*θ*) (i.e. population tuning curves) are ellipses with aspect ratios *k* := *ρ*_1_*/ρ*_2_ = 1, 3, and 5 respectively. The noise around each mean response point is i.i.d. standard Gaussian, so the total FI is exactly the length of the ellipse. We kept the length *S* (and thus the total FI) constant across the three codes by numerically solving the relevant equations composed of complete elliptic integrals of the second kind to get *ρ*_1_. Note that *r*(*θ*) is arclength-parameterized to ensure that FI is constant w.r.t. *θ*. It does not have an analytic form and has to be numerically computed by solving equations composed of the appropriate incomplete elliptic integrals of the second kind. To get the classification errors in the neurometric plot, we randomly selected 100 pairs of (*θ, θ*^′^) on the ellipse separated by Δ*θ*, sampled 10^4^ data points around each mean response location by adding i.i.d. standard Gaussian noise, and classified them according to their distance to the two means (which is the optimal classifier in this context). MI is estimated using Monte Carlo integration.

#### Tuning curves and selectivity

Fig. 3a shows example tuning curves from the von Mises model and the SoC model. The von Mises model is given by Eq. (S15), with the parameters chosen such that it matches the SoC model in FI and average maximum firing rate (Supplementary Information A.4). The SoC is generated by randomly sampling in ℝ^*N*^ the center and two vectors spanning the 2-dimensional subspace containing the circle. The selectivity of neuron *i* is defined to be the modulus of the complex number *z*_*i*_, where

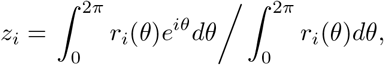

consistent with previous work [26, 27, 103].

#### Classification errors of tuning models

To compute the neurometric curve in Fig. 3b, we first generated population tuning curves with population size *N* = 100. (All simulations have *N* = 100 unless otherwise stated.) Again, the parameters of the von Mises model are chosen to match the FI and maximum firing rate of SoC (Supplementary Information A.4). For the nonnegative SoC, the circle *y*(*θ*) is sampled such that min_*θ*_ *y*_*i*_(*θ*) = 0. For the SoC with sign ambiguity, we allow *y*_*i*_(*θ*) to have negative values and set *β* = 4 (defined in Supplementary Information A.3), which leads to around 1/4 of neurons having bimodal tuning curves, consistent with neural data (Fig. 4d). Then, the classification errors are computed by sampling 100 pairs of (*θ, θ*^′^) separated by Δ*θ*, generating 2 × 10^4^ spike count vectors for each mean response according to the noise model (independent Poisson in the case of Fig. 3b), and classifying them using the likelihood ratio classifier.

For Fig. 3d, the parameter *σ* controls the strength of the gain modulation to the neurons’ firing rates (Eq. (3)). Each *σ* leads to a different optimal tuning curve (Supplementary Information A.5). During simulations, we sampled spike count data using one parameter (“*σ* of Gamma-Poisson noise”) and generated tuning curves using another (“*σ* of tuning curves”) while keeping the total FI constant by changing the radius of the circle *y*(*θ*). The plotted classification error is at Δ*θ* = *π/*2. To keep the comparison fair, we assume here that the decoder does not have access to the generating noise model and has to train a linear classifier based on 10^3^ data points from each *θ*. The classification errors are 5-fold cross-validated. Similarly for Fig. 6, different realizations of the connectivity matrix *W* create different variability structure (Eq. (S23)), which in turn produces different optimal tuning curves (Eq. (S24)). Again, we could either match or mismatch them when calculating classification errors while keeping FI constant, producing Fig. 6c.

#### Numerical optimization under population firing rate constraint

In Fig. 7, we numerically optimized population tuning curves under not just the length constraint, but also constant population firing rate at each *θ* and the monotonicity constraint Eq. (S26). The monotonicity constraint prevents degenerate behavior such as self-intersection, even when the length is unconstrained. We parameterized the tuning curves in *y* coordinates in the Fourier basis up to the 10th mode and used the fmincon function of MATLAB to solve the constrained optimization problem. The neurometric curves in Fig. 7c was computed by numerically integrating Eq. (2). An alternative approach is to discretize the curve in *θ* space as in [45], but the Fourier approach is more efficient as it evaluates the cost function with higher precision over a smaller number of variables.

### Data analysis

#### Tuning curves

The tuning curves used in Fig. 4 were calculated using the same method as the original paper [34] and the same criteria was used to identify HD cells among the recorded units. Each tuning curve was fitted to the sum of two von Mises curves, yielding a 6-dimensional nonlinear least-squares problem that was solved using MATLAB’s lsqnonlin function. The phase difference between the two peaks was constrained to be at least *π/*8 to avoid spurious bimodal fits that result from splitting single peaks in two. To rule out tuning curves that have three or more peaks, we only used those that are well fitted by the sum of two von Mises model (with *R*^2^ *>* 0.9) in Fig. 4d, which account for more than 80% of all HD cells.

The SoC code has 4 linear dimensions because squaring a (horizontally and vertically shifted) cosine function yields a linear sum of sin *θ*, cos *θ*, sin^2^ *θ*, cos^2^ *θ* and sin *θ* cos *θ*, which can be rear-ranged into a sum of sin *θ*, cos *θ*, sin 2*θ*, and cos 2*θ*. At the population level, this suggests that we can write

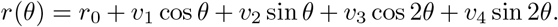

Then *r*(*θ*) is contained in a 4-dimensional affine subspace spanned by *v*_1_, *v*_2_, *v*_3_, *v*_4_ ∈ ℝ^*N*^ .

To get the *y* coordinates of tuning curves in Fig. 4f, we need to take the square root of the firing rates *r*_*i*_(*θ*) and assign the appropriate sign, which could be either positive or negative. This is achieved by fitting the SoC model (Eqs. (S10) and (S7)) to each neuron’s tuning curve and using the sign of the fitted *y*_*i*_(*θ*). More sophisticated models like the Gamma-Poisson model can be used to improve the variance explained by the fitting, but drastically increase the number of fitted variables, so we did not adopt them in order to avoid overfitting.

#### Variability structure of neurons

Fig. 5 relates two features of neuronal response in the data: the “super-Poissonness” *σ*_var_ and “tuning sharpness” *σ*_tun_. We first binned the HDs into 36 bins (i.e. 10^°^ bins) and then for each 100 ms time window counted the number of spikes as well as which bin the mouse’s HD falls into in this window on average. This yields a sampled distribution of spike counts for each HD bin. We then fitted the negative binomial distribution (Eq. (S19)) to this list of spike counts where the averages are allowed to vary across HDs but the variance of the gain modulation 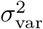 is held constant, similar to the procedure in [38]. This produces the super-Poissonness of a neuron. For the tuning sharpness, we fit Eqs. (S10) and (S21) to the tuning curves to obtain *σ*_tun_. Note that the fitted model fixes the peaks of bimodal tuning curves to be separated by *π* due to it being a transformation of cosine, but as seen in Fig. 4d, the peak phase difference of some bimodal tuning curves are not exactly *π*, which may produce spurious effects on *σ*_tun_. We therefore only use unimodal tuning curves in Fig. 5c, d, but the result holds qualitatively if we use all tuning curves. Since super-Poissonness and tuning sharpness are predicted to be the same in the model, they are denoted with the same notation *σ* in the theoretical derivations, but we use different notations *σ*_var_ and *σ*_tun_ here to emphasize that they are fitted independently from different aspects of the neuronal activity data. We normalized the fitted values among the recorded population because spike trains are fundamentally point processes and the statistics of spike counts heavily depend on the time window used [104]. It is impossible to determine the temporal structure of downstream decoders, so our theoretical prediction on the agreement between *σ*_var_ and *σ*_tun_ can only be expected to be qualitative. That is, it is relative between neurons in the population. Fig. S2 is produced using the same procedure but for the monkey V1 data in [30].

One potential confounding factor is the effect of differential correlations [16]. For example, in a ring attractor model of the maintenance of HD information [99, 100], there is a dynamic inclination to drift along the ring. Then neurons with a sharper tuning curve has a larger loading in this particular direction (i.e. *r*^′^(*θ*)), which may lead to larger variances if differential correlations dominate the population variability. To rule out this effect, we subtracted the component along the *r*^′^(*θ*) direction (which is estimated by the difference of the mean responses to neighboring *θ*’s) from the centered spike count data before estimating *σ*_var_. The results in Fig. 5c, d are virtually the same with or without this procedure.

## Data availability

The mouse HD cell dataset used in this paper was previously published [34] and is publicly available at https://doi.org/10.6084/m9.figshare.24921252. The monkey V1 dataset used in this paper was previously published [30] and is publicly available at https://crcns.org/data-sets/vc/pvc-11/about.

## Code availability

Simulation and data analysis codes are provided at https://github.com/brain-math/globally-optimal-neural-code.

## Acknowledgments

We thank E. Simoncelli and L. Snyder for insightful discussions. Mouse cartoon in Fig. 4a downloaded from https://doi.org/10.5281/zenodo.3925997.

## Supplementary Information

### A Mathematical derivations

#### A.1 Problem statement

Suppose a population of *N* neurons represents a variable *θ* lying on some manifold *M* with its activity *x* ∈ ℝ^*N*^ . While this variable is not limited to sensory stimuli but may be cognitive or motor in nature, we refer to it as the “stimulus” in this paper for convenience.

Due to trial-to-trial variability in the response, the neural code is a mapping

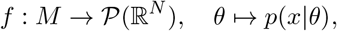

where *P*(ℝ^*N*^) is the space of neural response distributions. The quality of a population code is measured by a cost function *L*(*f*) and a code is said to be optimal if it minimizes *L*(*f*). Different computational goals correspond to different cost functions. In this paper, we focus on one-dimensional circular variables, namely *M* = *S*^1^ = [0, 2*π*].

In the context of promoting discriminability, previous works often use Fisher information (FI) [14, 16, 17, 105] to the measure the quality of the neural code. It is defined as

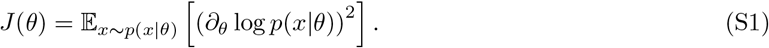

However, as discussed in the main text, FI is fundamentally local and does not capture the global structure of the code. Mutual information is a global measure, but is difficult to study analytically. We therefore aim to develop a cost function that is both global and amenable to mathematical analysis that allows us to discern representations that have the same FI but different global properties. Here, by the same FI we first mean that the total FI

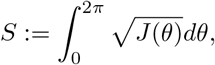

is held constant, as previous works have argued that the total FI sets the overall available coding resource or model complexity [9, 21, 29, 43–45]. We additionally assume for now that the prior distribution of *θ* is uniform over [0, 2*π*]. Efficient coding principles then suggest that the optimal code should satisfy 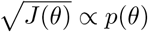 [21, 29], so we require FI to be constant over *θ*.

In sensory discrimination tasks, classification errors are often hypothesized to arise from the overlap between the distributions *p*(*x* | *θ*) and *p*(*x* | *θ*^′^) in response to the pair of stimuli (*θ, θ*^′^) between which downstream neurons need to distinguish. Let us start by selecting a measure of how close two distributions are in neural activity space, i.e. a distance function in *P*(ℝ^*N*^). There are many statistical distances to choose from. The total variation distance and the Hellinger distance are directly related to the classification error between distributions [106], but are hard to analyze for our purposes as will be seen later. We adopt instead the Fisher-Rao distance from the field of information geometry [47, 48]. (Note that the Fisher-Rao distance can be thought of as the inherited metric on a parameterized submanifold of the full space of probability distributions equipped with the Bhattacharya angle as the distance, which is in turn inherited from the Hellinger distance on the space of all measures [107]. Interestingly, the Fisher-Rao distance can be written in terms of the Hellinger distance for the case of independent Poisson variability [107].) It is based on the observation that for a family of distributions fully described by a set of parameters, the FI matrix with respect to those parameters is positive definite and constitute a Riemannian metric. Then this space of distributions becomes a Riemannian manifold with those parameters as the coordinate system. In our case, notice that the neural response distribution is often determined by the mean response E[*x*] ∈ ℝ^*N*^ . That is, the map

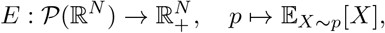

where 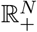 is the nonnegative orthant of ℝ^*N*^, is a bijection. We then denote *p*_*r*_ := *E*^−1^(*r*). For example, the vector of firing rates is sufficient to completely specify the joint distribution of a neural population’s spike counts if we assume that they are independent Poisson neurons. In particular, (with a slight abuse of notation) this bijection allows us to pull back the map *f* : *M* → *P*(ℝ^*N*^) to 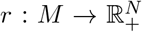, which is the population tuning curve. Following information geometry as stated above, we give ℝ^*N*^ the FI metric:

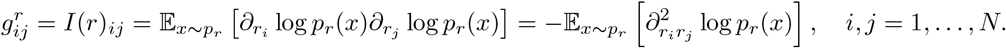

We denote this Riemannian manifold by *K* and the geodesic distance on it by *d*(·, ·). It is important to emphasize that the FI matrix used here is with respect to the mean response *r* ∈ ℝ^*N*^ and the defined metric structure exists for the full space of neural response distributions independently of the stimulus *θ* being encoded, in contrast to Eq. (S1).

Naively, we would want to maximize the distance between the neural response distributions and the cost function would simply be the average of the negative distance −*d*(*r*(*θ*), *r*(*θ*^′^)) across all pairs (*θ, θ*^′^). However, this fails to capture the fact that increasing the distance between two distributions beyond a certain point should yield no further benefit, as there will be little chance of mistaking one for the other after that point. This suggests that the cost contributed by each stimulus pair (*θ, θ*^′^) should be a function *h* of the distance *d* that decreases to 0 as *d* grows large enough. Ideally *h* would be the classification error, but unfortunately the classification error is not a function of the Fisher-Rao distance. However, this is approximately the case for small distances, which is sufficient for our purposes since only then is the classification error large and behaviorally relevant.

Let us derive this approximation. The classification error for a decoder based on the likelihood ratio is

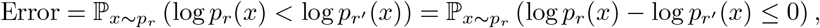

where we have assumed symmetry for simplicity. For independent or weakly correlated neurons, the random variable 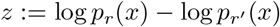 is roughly Gaussian. Then we have:

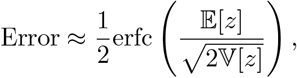

where erfc is the complementary error function. Since

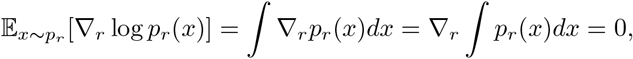

we have, using Taylor expansion,

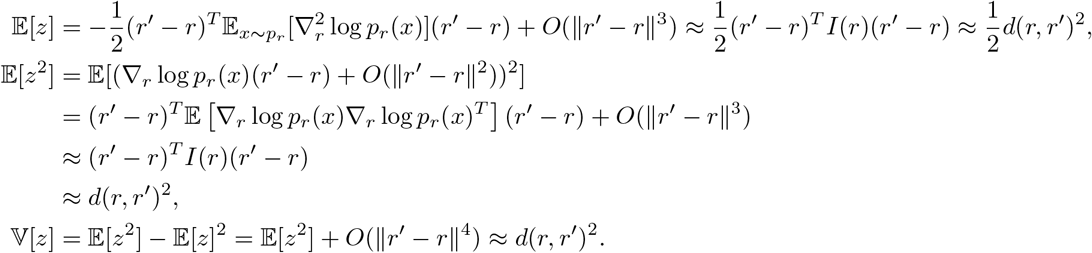

Therefore

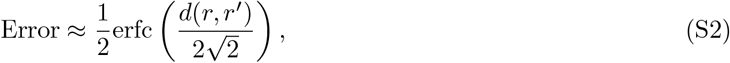

and we can define

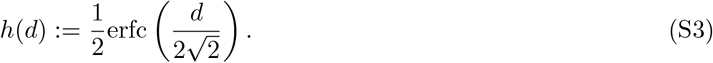

This is consistent with results from previous work [14] in the sense that if the neural code passes through *r* and *r*^′^ with *r*(*θ*) = *r* and *r*(*θ* + Δ*θ*) = *r*^′^, then by the chain rule

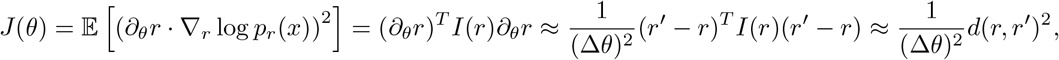

and we recover the formula from [14]

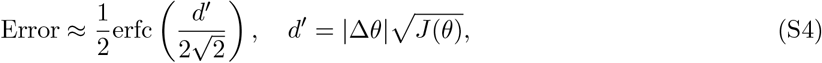

where *d*^′^ (“d-prime”) is the discriminability index. The key difference is, Eq. (S2) describes the classification error between any pair of distributions with mean response *r* and *r*^′^ regardless of what stimulus the population is encoding (or lack thereof). This captures the relationship between points on the population tuning curve *r*(*θ*) even if they are separated by a large Δ*θ*, setting the stage for understanding the global structure of the neural representation, whereas the result Eq. (S4) from [14] is fundamentally local as it only applies to small Δ*θ*’s.

With the above preparations, we can finally define our cost function as the (approximate) mean classification error:

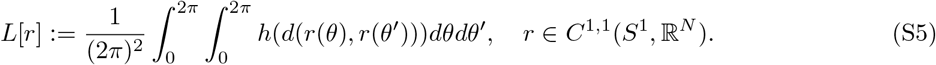

Here *C*^1,1^(*S*^1^, ℝ^*N*^) is the (infinite-dimensional) function space of smooth closed curves in ℝ^*N*^, making this a variational problem. The double integral runs through all pairs of (*θ, θ*^′^) regardless of the size of Δ*θ* = *θ*^′^ −*θ*, making this a nonlocal variational problem [74].

Recall that our aim is to use *L*[*r*] to distinguish between population codes with the same FI. Under the metric *g*^*r*^, the FI (as defined in Eq. (S1)) of the code *r*(*θ*) is the square of the speed of the curve, i.e., 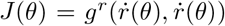, and the length of the curve is the total FI [21, 29]. Then the statement of our problem is completed by requiring the closed curve *r*(*θ*) to have constant speed with a given length *S* in Eq. (S5). One can view a constraint on FI (i.e., the speed of the curve *r*(*θ*)) as a regularizing condition on the optimization problem, as it prevents the curve from expanding unboundedly when trying to the separate the points on the curve from each other as much as possible.

#### A.2 Solving the nonlocal variational problem for independent Poisson neurons

We solve in this section the nonlocal variational problem Eq. (S5) under the assumption that neurons have independent Poisson variability.

Although the FI of the Poisson distribution is a standard result [14], let us nevertheless calculate it here for illustrative purposes. The neural response distribution family is

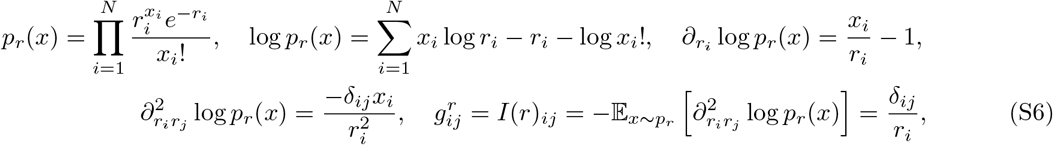

where *δ*_*ij*_ is the Kronecker delta. Then *g*^*r*^ is a diagonal matrix where each *i*^th^ entry on the diagonal only depends on the *i*^th^ coordinate, making the metric Euclidean. This is true for any noise model where neurons are independent (in the sense that 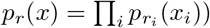, as the statistical manifold would then be the product of *N* manifolds [48]. To make this explicit, consider the coordinate transform:

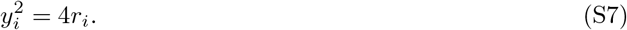

Then the metric in *y* coordinate is

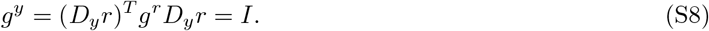

More generally, one can solve Eq. (S8) whenever the FI metric *g*^*r*^ is Euclidean to get the alternative coordinate system *y*, as we will do below in sections A.5 and A.6. Then the geodesic distance in *y* coordinate is explicitly the Euclidean distance, that is

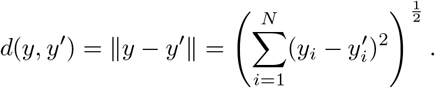

The optimization problem then becomes min *L*[*y*] where

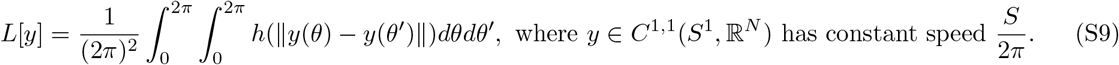

This has been solved by existing work in knot theory [46]. We restate their theorem here:

##### Theorem 1

(Theorem 2 in [46]). *For a function g*(*d*_1_, *d*_2_) *from* ℝ^2^ *to* ℝ *that is convex and decreasing in d*^2^ *for d*^2^ ∈ (0, *d*^2^] *and d*_2_ ∈ (0, *π*), *the energy*

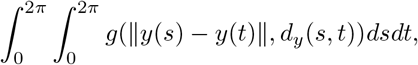

*is uniquely minimized among closed unit-speed curves of length* 2*π by the round unit circle. Here d*_*y*_(*s, t*) *is the distance along the curve y between the points y*(*s*) *and y*(*t*).

In particular, this theorem holds if *g*(*d*_1_, *d*_2_) is constant over *d*_2_, and being convex in *d*_1_ is sufficient for it to be convex in 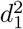. It is easy to show that the *h* defined in Eq. (S3) is convex and decreasing. Therefore as a direct corollary, we see that our variational problem Eq. (S9) is solved by round circles of length *S*, that is

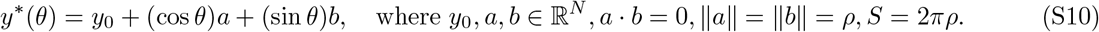

This can be transformed back to firing rate coordinates using Eq. (S7) to get the population tuning curve *r*(*θ*). Since the transformation for independent Poisson neurons is the element-wise square, we call this optimal code the “square-of-circle” code.

In fact, the lemma used to derive the above theorem implies a stronger result. Let us restate it here:

##### Theorem 2

(Theorem 8 in [46]). *Let y be a closed, unit-speed curve of length* 2*π in* ℝ^*N*^ . *For s* ∈ (0, 2*π*), *if* 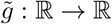 *is increasing and concave on* (0, *d*_*y*_(0, *s*)^2^], *then*

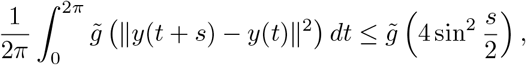

*and equality holds if and only if y is the unit circle*.

This theorem tells us that the code Eq. (S10) not only minimizes the average classification error, but is also optimal at every point of the neurometric curve. That is,

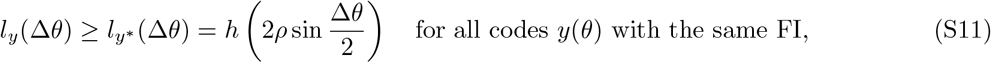

where

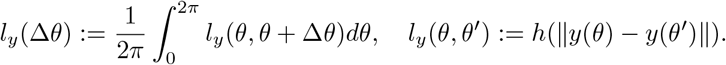

*l*_*y*_(Δ*θ*) is the average classification error for pairs of *θ*’s separated by Δ*θ*, i.e. the neurometric curve. In the other direction, *L*[*y*] is the average of *l*_*y*_(Δ*θ*) over Δ*θ*.

#### A.3 The impact of sign ambiguity

From this section onward all curves *y*(*θ*) in *y* coordinates we consider will be round circles in the form of Eq. (S10), and we omit the asterisk for convenience. We will also take a probabilistic approach and assume that the neurons’ tuning curves are sampled from some distribution, with assumptions on this distribution specified as we progress.

Our result in the previous section seems to suggest that *any* round circle *y*(*θ*) with the same length achieves the optimal performance in the binary classification task. However, this statement needs a slight modification to be strictly accurate: the elements of *y*(*θ*) have to be nonnegative for all *θ* for the resulting neural code to be optimal. The reason for this is that the map Eq. (S7) between *y* and *r* coordinates is only a bijection for each orthant of *y*. When there exists *i* such that *y*_*i*_ and 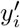 have different signs, we must have *d*(*r, r*^′^) *< d*(*y, y*^′^). More specifically,

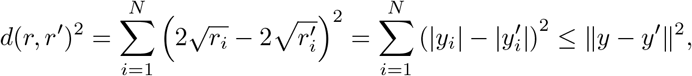

with equality if and only if *y* and *y*^′^ are in the same orthant (i.e. *y*_*i*_ and 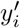 have the same sign for all *i*).

This implies that a round circle *y*(*θ*) is only optimal if it lies in one orthant, which for our purposes can be chosen to be the nonnegative orthant. Recall that the tuning curve *r*_*i*_(*θ*) is bimodal if and only if *y*_*i*_(*θ*) has both positive and negative values. Then the bimodal tuning curves observed in data (Fig. 4d) would imply that the neural code is suboptimal. However, we shall next show that this degradation of performance due to sign ambiguity is vanishingly small.

Let us calculate the classification error for the circle in Eq. (S10) allowing sign changes. Denote the element-wise absolute value of a vector *v* ∈ ℝ^*N*^ by 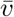. Then

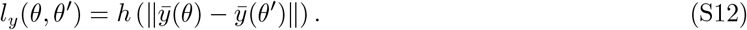

Without loss of generality, we may assume that *y* intersects coordinate hyperplanes at a finite number of points, and we call the transverse intersections zero-crossings. We also assume that each intersection is with only one hyperplane, since if not, an arbitrarily small perturbation can split that special crossing point into two generic ones. Then each coordinate hyperplane (i.e. neuron) is either never crossed or crossed twice.

Neuron *i*’s tuning curve can be written in two forms:

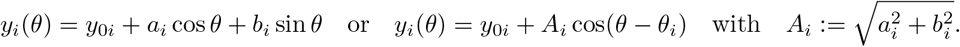

Then *θ*_*i*_ is the preferred stimulus of neuron *i*. If neuron *i* has two zero-crossings at *φ*_*i*_ and 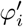 with 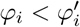 and 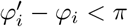 (modulo 2*π*), we have:

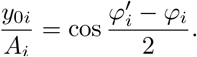

Notice that neuron *i* contributes to the sign ambiguity only if it has exactly one zero-crossing between *θ* and *θ* + Δ*θ*. Denoting the set of these neurons by *I*_*θ*_, we have:

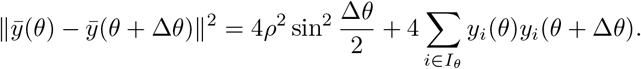

For convenience, define

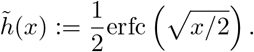

Then

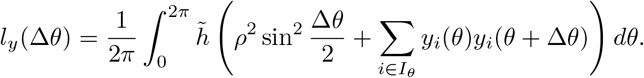

Let us assume that the preferred stimuli *θ*_*i*_’s are distributed uniformly in [0, 2*π*]. Then we can assume that *y*_0*i*_ ≥ 0, as a negative *y*_0*i*_ would have the same effect on the result if we change *θ*_*i*_ to *θ*_*i*_ + *π*. When *N* is large, *ρ*^2^ ∼ *O*(*N*) and a finite Δ*θ* would lead to |*I*_*θ*_| ∼ *O*(*N*). For large *N*, we can treat *θ* as a uniform random variable and make the approximation

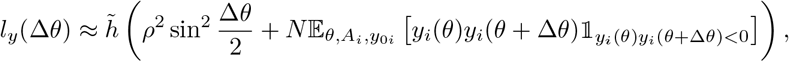

where 1_*X*_ is the indicator function of set *X*. Since 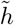 is convex, by Jensen’s inequality this approximation is in fact a lower bound.

Assuming the overall magnitude *A*_*i*_ and the relative parameter *x* := *y*_0*i*_*/A*_*i*_ are independent, we have

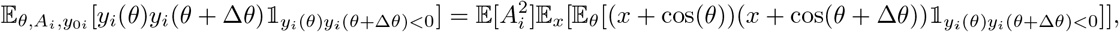

in which

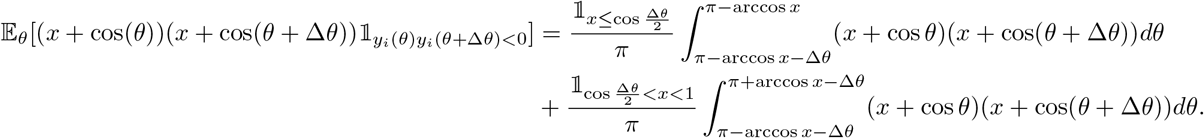

These integrals reduce to:

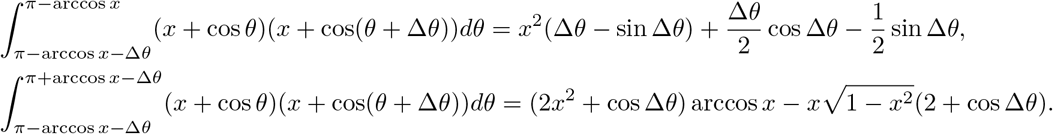

In total, we have:

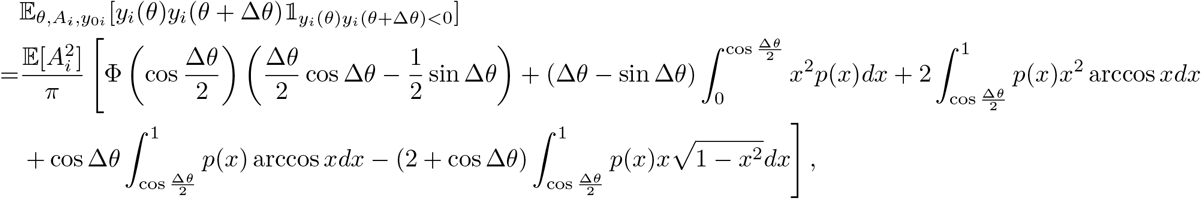

where *p* and Φ are the pdf and cdf of *x* = *y*_0*i*_*/A*_*i*_ respectively.

For concreteness, let us calculate the case of 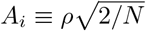 and *x* = *y*_0*i*_*/A*_*i*_ ∼ *U*(0, *β*), *β >* 1. Then

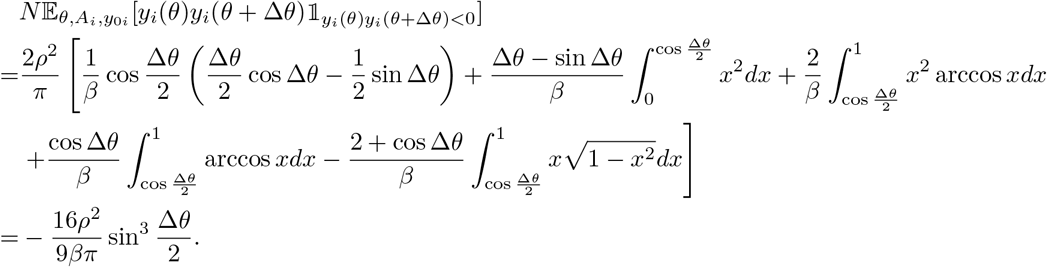

We arrive at

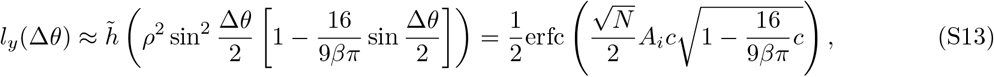

where *c* := sin(Δ*θ/*2). We argue that sign ambiguity has a very small effect on the classification error. To see this, let 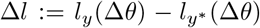 where *y* *(*θ*) now denotes the truly optimal code, i.e. a nonnegative circle. We have

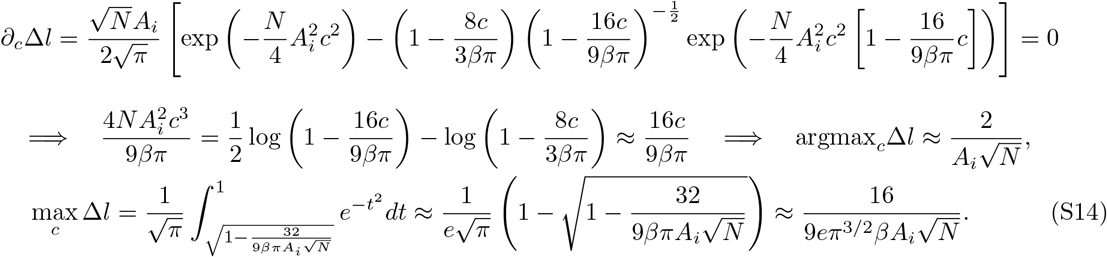

(Note that *A*_*i*_ is the dynamic range of neuron *i* and stays *O*(1) while the radius 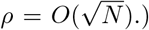 This is a uniform bound on the expected increase in classification error for models with sign ambiguity for all Δ*θ*’s. It is thus negligible for large *N* at any Δ*θ* where the error is finite. While this is a lower bound, we verified with numerical calculation of Eq. (S12) that it is a good approximation for large *N* (Fig. S1).

What this result says is that the mesoscopic structure of the circle (i.e. the behavior of *y*(*θ*) in each orthant) dominates the final performance of the neural code, and that the increase in classification error due to sign ambiguity is vanishingly small for large population sizes. The bimodal tuning curve observed in data thus does not imply a large deviation from optimality. Also note that a code with sign ambiguity has smaller firing rates than exactly optimal codes with the same FI. The latter achieves minimum firing rates when *y*_0*i*_ = *A*_*i*_ (i.e. *x*_*i*_ = 1), while the former can reduce the firing rates by allowing |*x*_*i*_| *<* 1. Then bimodal tuning curves can be viewed as a trade-off between discriminability and energy consumption.

#### A.4 The von Mises model

We used the classical von Mises model [41, 45, 55–57] as a baseline comparison for our square-of-circle code. In this section, we calculate its performance analytically. Throughout this section, we will repeatedly use the following properties of modified Bessel functions of the first kind:

1. *I*_0_(*κ*) − *I*_2_(*κ*) = 2*I*_1_(*κ*)*/κ*;
2. 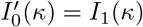
3. 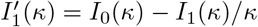

The von Mises model is

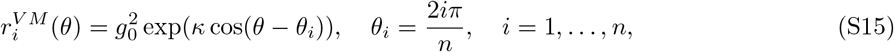

with independent Poisson noise. The parameter *κ* controls the width of the tuning curves, with a larger *κ* leading to narrower tuning. One can view this as using an alternative nonlinearity (exponential instead of the square) on the cosines, making it an “exponential-of-circle code”. In *y* coordinates where the Fisher-Rao distance is Euclidean, this curve is

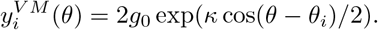

Then the FI is

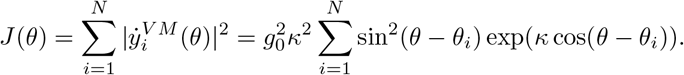

For large *N* and uniform tiling of preferred orientation *θ*_*i*_, *J*(*θ*) would be virtually constant w.r.t. *θ* and we can calculate the analytic expression for the length of the curve to be

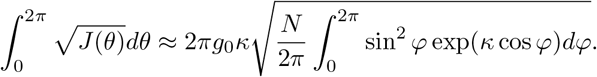

Then fixing the total FI, i.e. fixing the length at *S*, can be achieved by setting

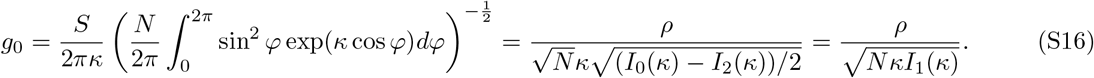

The neurometric curve is

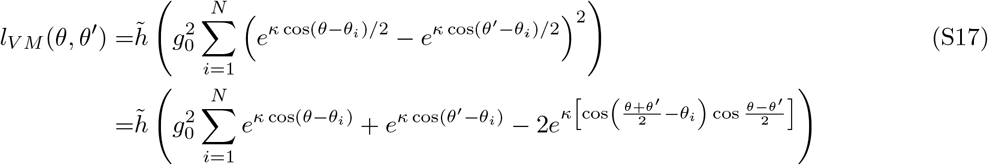

It is difficult to simplify further, but we can look at its approximate behavior by taking the continuous limit of large *N* to get

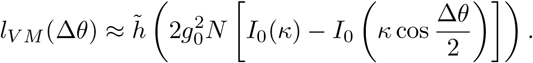

Again, by Jensen’s inequality, this is in fact a lower bound. Substituting the normalizing scaling in Eq. (S16) to keep the total length constant (*S* = 2*πρ*), we have

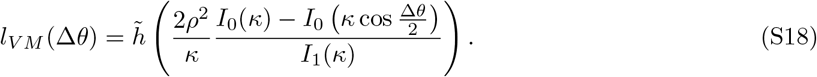

We will next show that *l*_*V M*_ (Δ*θ*) is monotonically increasing with *κ* for each Δ*θ*. Let 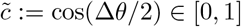. We only need to show that

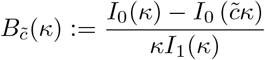

is monotonically decreasing. Taking its derivative gives us

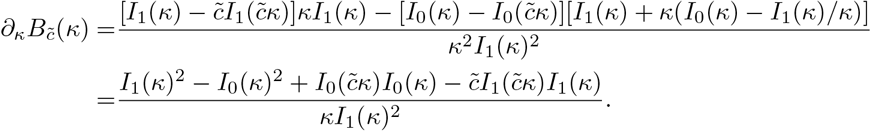

Then we only need to prove that

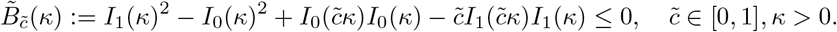

Notice that when 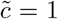 we have 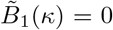, so we will be done if we can prove that 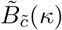 monotonically increases with 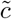 for all *κ >* 0. Let us take the derivative

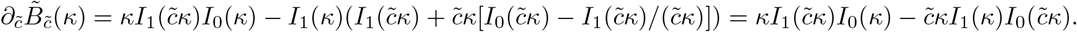

It has been previously shown that the function *xI*_0_(*x*)*/I*_1_(*x*) is monotonically increasing in *x* [108]. Then because 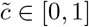, we have

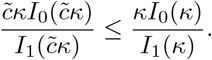

Rearranging this give us 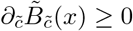, which is the desired result. Then *l*_*VM*_ (Δ*θ*) is monotonically increasing with *κ* for each Δ*θ*. This says that when FI is fixed, the broader the tuning curve the better the classification performance. Interestingly, this is the opposite to the case where the maximum firing rate is kept fixed and FI is allowed to vary. There, previous work has shown that FI is larger for sharper tuning curves in 1D [15]. This demonstrates the opposite effects of tuning width on local and global discriminability, and this trade-off has been shown numerically in previous work [22].

Let us then calculate the limit of *κ* → 0 to get a lower bound:

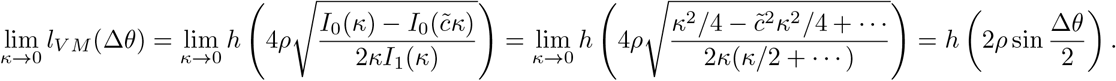

We see that it converges to 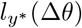, recapitulating the optimality of the square-of-circle code. This can also be obtained by taking *κ* → 0 directly in Eq. (S17), suggesting that the von Mises model’s performance (and not just a lower bound) approaches that of the square-of-circle code for small *κ*. However, *κ* cannot be too small because the maximum firing rate of a neuron is

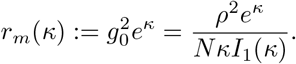

Clearly, *r*_*m*_ → ∞ as *κ* → 0. To make a fair comparison, we therefore need to match the (average) maximum firing rates of the models by setting

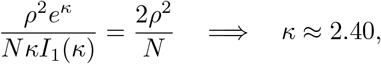

where we have assumed *y*_0*i*_ = *A*_*i*_ (i.e. *x*_*i*_ = 1) which minimizes the firing rate among all nonnegative SoC codes.

#### A.5 Gamma-modulated Poisson noise

In the next few sections we consider noise models other than independent Poisson distributed spike counts. Consider a multiplicative gain term *G* that modulates the firing rate [38]. That is, *x*_*i*_ ∼ Poiss(*r*_*i*_*G*_*i*_), where *G*_*i*_ follows a Gamma distribution with mean 1 and variance 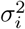 that is independent across neurons and does not depend on *r*. Under this model (and marginalizing over *G*_*i*_), *x*_*i*_ follows a negative binomial distribution 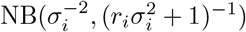 and

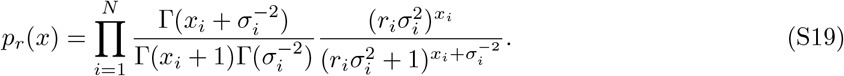

The variance is

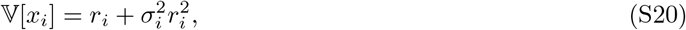

and the FI (w.r.t. *r*) is

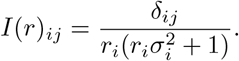

Then we can solve Eq. (S8) to get the appropriate transformation

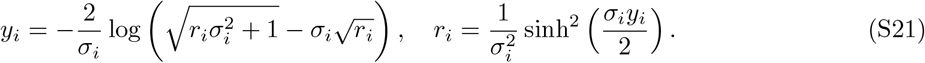

Qualitatively, *r*_*i*_ is an even function of *y*_*i*_ that increases monotonically and supralinearly with |*y*_*i*_|, with a larger *σ*_*i*_ leading to a more supralinear relation (Fig. 3c). Notice that we can recover the squaring transform from independent Poisson by letting *σ*_*i*_ → 0. This transformation is exactly the same as what was used in [62] (see Eqs. (7) and (8) of that paper), but they used this procedure for a different purpose and did so heuristically without establishing a Riemannian metric structure as we did here.

In Fig. 3d, we demonstrate the dependence of the optimal tuning curve (Eqs. S10 and S21) on the noise structure (Eq. (S19)) and the importance of matching the former with the latter. For simplicity, we consider the case where all neurons have the same *σ*_*i*_ parameter in their variability, that is *σ*_*i*_ = *σ* for all *i*. Write the noise model (Eq. (S19)) as *p*^*σ*^ : *r* ↦ 1 *p*_*r*_ and the corresponding transformation between *r* and *y* coordinates (Eq. (S21)) as *r*^*σ*^ : *y* ↦ 1*r*. For each *r*^*σ*^, we can get a population tuning curve by using it to transform a circle *y*(*θ*), which is optimal for the noise model *p*^*σ*^ but suboptimal for 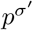 if *σ*^′^≠ *σ*.

In Fig. 5, we allow *σ*_*i*_ to be different for different neurons and examine how the variability of each neuron relates to its tuning curve shape *within the same* population code that is hypothesized to be optimal. This is in contrast to Fig. 3d where we compare performance *across different* codes indexed by *σ*.

### A.6 Noise correlations from a linearized recurrent network

The noise models in previous sections all have independent neurons. To illustrate that our framework can also incorporate noise correlations, we consider a covariance structure that is generated by a recurrent circuit. Specifically, consider

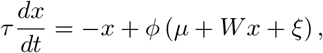

where *W* is the recurrent connectivity matrix, *µ* is the mean input vector, *ξ* ∼ *N*(0, *C*_0_) is trial-to-trial noise, and *ϕ* is a transfer function that applies element-wise. Assume that the network evolves to a steady state in each trial. Then

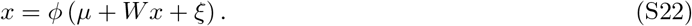

When the noise *ξ* is small, we can linearize the above equation to get

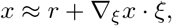

where *r* is mean response, which is the steady state when there is no noise, i.e.

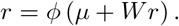

Differentiating Eq. (S22) gives us

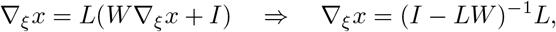

where *L* := diag(*l*_1_, …, *l*_*N*_) is the diagonal matrix of neuronal gains

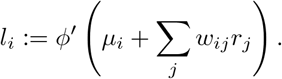

Then

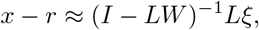

and the covariance matrix *C* is

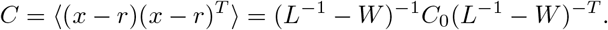

This is a standard result for population covariability in a linearizable recurrent network [63–65].

Experiments have shown that the transfer function of neurons can be well described by a threshold-quadratic law [109, 110]. This leads to a relationship between the neuronal gain and the firing rate: 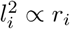. Writing *R* := diag(*r*_1_, …, *r*_*N*_), we thus have [65]

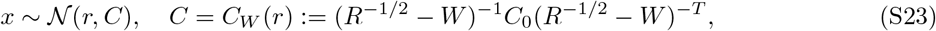

where we absorbed the constant in the 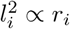 relation into *C*_0_ and *W* . The notation *C* = *C*_*W*_ (*r*) highlights the fact that the covariance matrix *C* is a function of the mean response *r*, and that this relationship depends on the recurrent connectivity *W* . Then there is again a bijection between the mean response *r* and the full neural response distribution, and we can proceed as we did before to establish a Riemannian metric on the mean response manifold. For simplicity, we use the linear FI in this section [66]. Then the Riemannian metric is *g*^*r*^ = *C*^−1^. We claim that the transformation

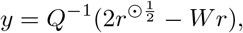

where *C*_0_ = *QQ*^*T*^ and *v*^⊙*p*^ is the element-wise *p*-power of *v*, makes the metric explicitly Euclidean. Indeed,

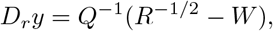

so

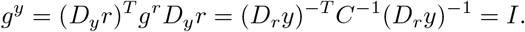

That is, our proposed transformation solves Eq. (S8). Notice that setting *W* = 0 recovers the transformation Eq. (S7) for independent Poisson neurons. Then the *linear* FI of an uncoupled population here equals the *full* FI of independent Poisson.

The optimal code in *y* coordinates is again the round circle Eq. (S10). Given *y*, we can solve for *r* numerically using the system of quadratic equations

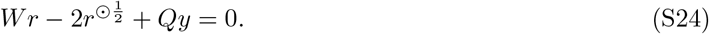

In general, a system of *N* quadratic equations can have at most 2^*N*^ solutions. (Simply consider each equation as having 2 roots and combine them.) Here we have additional constraints that makes the solution generically unique. For the network to be linearizable such that Eq. (S23) holds, we need to have spectral radius *ρ*(*LW*) *<* 1. Then the set of admissible solutions for *l* is

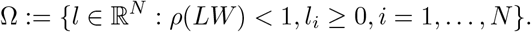

When *W* is i.i.d. Gaussian with *w*_*ij*_ ∼ *N*(0, *g*^2^*/N*), we have for large *N* [111]

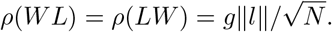

This says that Ω is the nonnegative orthant of a ball, and is in particular a convex set. Suppose there are two solutions to Eq. (S24), 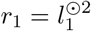 and 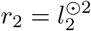 with *L*_*i*_ := diag(*l*_*i*_), *i* = 1, 2. Then

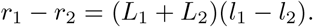

Subtracting the two equations then gives us

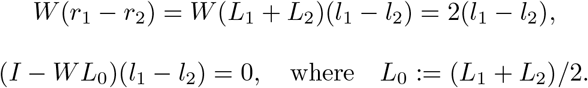

If *l*_1_ ≠ *l*_2_, then *ρ*(*WL*_0_) ≥ 1 and *l*_0_ := (*l*_1_ + *l*_2_)*/*2 ∉ Ω, which contradicts the convexity of Ω. Therefore the solution to Eq. (S24) is usually unique when *W* is a large i.i.d. Gaussian matrix and the recurrent circuit is linearizable.

#### A.7 Globally shared gain modulation produces hyperbolic geometry

In section A.5, we considered independent gain modulations for Poisson neurons. However, the separation of variability into “point-process” and “gain” components also partitions the correlations [38]. The simplest model in this setting is to have a global gain that is shared among all neurons. Let us see what the optimal code looks like in this case.

Let *G* again follow a Gamma distribution with mean 1 and variance *σ*^2^. That is, *G* ∼ Gamma(*σ*^−2^, *σ*^2^) under the scale characterization. The response of neuron *i* is *x*_*i*_ ∼ Poiss(*r*_*i*_*G*). Then the joint PMF is

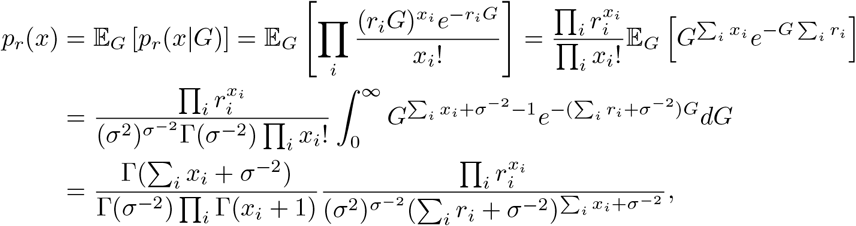

and

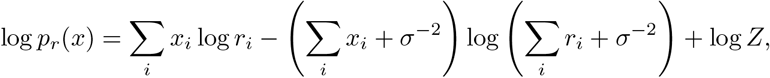

where *Z* does not depend on *r*. For *i* ≠ *j*,

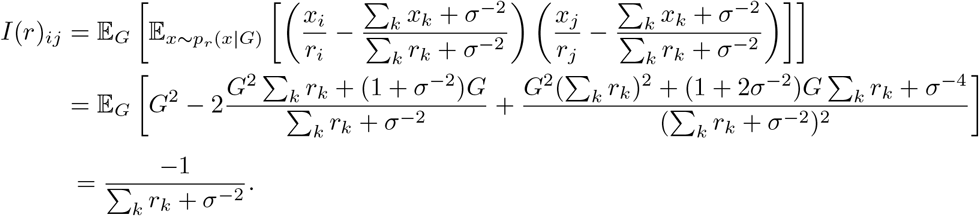

In addition,

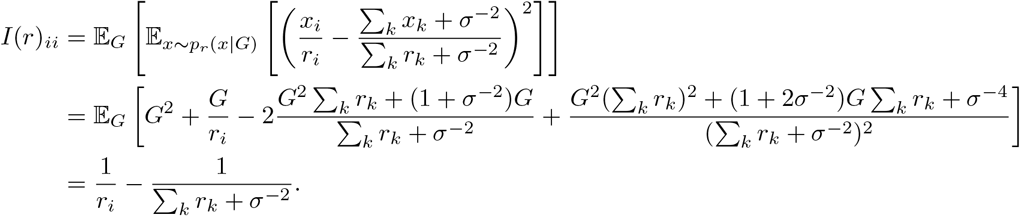

Taken together, we have

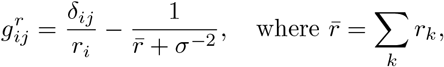

or in matrix form

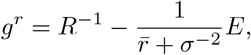

where *E* is a matrix of 1’s. This in fact makes the Riemannian manifold *K* the hyperbolic space, which has constant negative curvature. To see this, consider the hyperboloid model of hyperbolic space [112] with curvature −*σ*^2^*/*4:

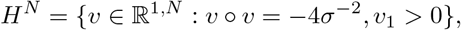

where ° is the Lorentzian inner product 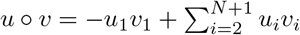. We claim that

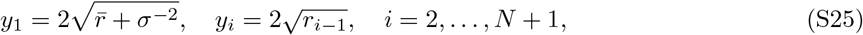

maps 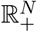 to *H*^*N*^ isometrically. Indeed, first notice that in the above formula

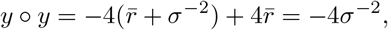

so the image of the mapping indeed lies in 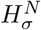. Additionally,

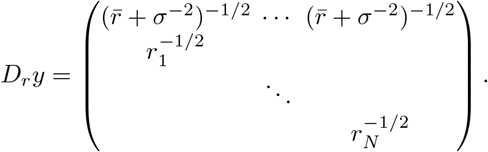

Then setting *g*^*y*^ = (−1, 1, …, 1) gives us

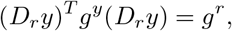

proving the isometry.

In the following, we shall assume *σ* = 2 for simplicity. Theorem 1, on which our result is based, does not directly apply to non-Euclidean manifolds. However, we can recover our result in the hyperbolic space case with a mild monotonicity requirement. Specifically, we assume that the representation *r*(*θ*) preserves the semantic structure of *S*^1^ in the sense that

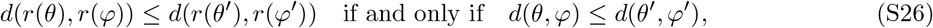

where

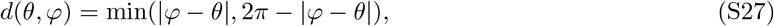

is the distance on *S*^1^. This essentially ensures that the neurometric curve is well-defined and monotonically decreasing. In particular, it implies that *d*(*r*(*θ*), *r*(*φ*)) is a function of *d*(*θ, φ*). (A weaker constraint that only requires *d*(*r*(*θ*), *r*(*φ*)) to be a function of *d*(*θ, φ*) is the “translation invariance” given in [45]. Their optimizations indeed yield neurometric curves that are not monotonic, which is not behaviorally realistic.) In words, this condition reads “stimuli that are close semantically are close in their representations”. In Euclidean space, this requirement is automatically satisfied by the optimal code in Eq. (S10). In hyperbolic space, this implies that for any *t* ∈ [0, 2*π*),

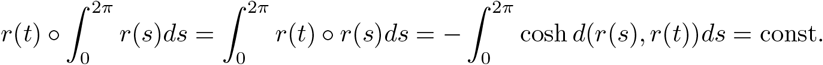

This says that the curve *r*(*t*) is contained in an affine hyperplane *P* that is Lorentzian orthogonal to 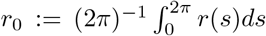 and contains *r* . Since *r* is time-like and *P* is space-like, the Lorentzian inner product is Euclidean in *P* . Then we can again use theorem 1 to prove that the minimizers of *L*[*y*] under the monotonicity constraint are round circles in *H*^*N*^ . Interestingly, simulations show that circles are the minimizer of *L*[*y*] even without this constraint, but we have yet to obtain a proof.

The easiest way to describe what a “circle” is in *H*^*N*^ is to take the center of the circle to be 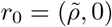 where 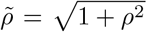. Then 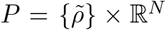 in which the horizontal hyperplane ℝ^*N*^ is explicitly Euclidean and constructing a circle there is straightforward. (This corresponds to holding the population firing rate 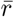 constant across *θ*.) We can then transform it with a hyperbolic isometry (which is a linear map in *R*^1,*N*^) to get a general circle in *H*^*N*^ . This would produce an ellipse in ℝ^*N*^ (as the coordinate subspace of ℝ^1,*N*^ that discards the first coordinate). Then Eq. (S25) tells us that the resulting population tuning curve in *r* coordinates is a “square-of-ellipse”, which has no features that are qualitatively different from the independent Poisson model and thus difficult to test in neural data.

#### A.8 Constraint on population firing rate produces spherical geometry

Let us return to the independent Poisson model where the appropriate coordinate transformation is Eq. (S7). If we require that the population firing rate 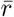 to be constant for all *θ* following some energy consumption considerations, then the curve *y*(*θ*) is constrained to lie on the sphere 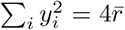. Since there is a one-to-one mapping between the geodesic distance on the sphere (which are great arcs) and the Euclidean distance in the ambient space, placing this constraint is equivalent to letting the manifold *K* be the (nonnegative orthant of the) sphere, which has constant positive curvature.

When the length of the curve *S* is smaller than that of the largest positive circle (LPC), which has length 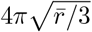 (i.e. 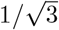 times that of the great circle), it is clear that the solution to the variational problem is again the round circle. (Here the LPC has the critical length instead of the great circle because the issue of sign ambiguity is very prominent due to the constraining sphere being centered at the origin. Intuitively, every point on the great circle would be confused with its opposite point when mapped back to *r* coordinates.) When 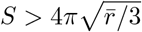, however, there no longer exists a positive round circle in the sphere of length *S*, and the situation is much more complicated. There is no equivalent version of theorem 1 on the sphere, even with the monotonicity requirement defined in the previous section (Eq. (S26)).

Furthermore, in this scenario it is also interesting to remove the constraint on the length of the curve and only constrain the population firing rate to be constant, as the compactness of the sphere already acts as an effective regularization. A variant of this problem has been discussed and partially solved by previous work [113], but only for *N* = 3. In addition, the solutions they obtained do not respect monotonicity (Eq. (S26)), as the compactness of the space forces the curve to “circle back” when its length is larger than that of the great circle. Therefore, we turn to numerical optimization in Fig. 7.

A previous work examined a similar optimization problem [45]. They assume that the *l*^2^ norm of the firing rate vector is constant instead of its sum (i.e. *l*^1^ norm), but they use isotropic Gaussian noise as opposed to Poisson noise, so the distance on their sphere is the same as our sphere in *y* coordinates. There are three crucial differences between their formulation and ours. First, they optimize the Coulomb energy (i.e. *h*(*d*) = 1*/d*) instead of the average classification error Eq. (S5). Second, they only assume translational invariance instead of the monotonicity constraint Eq. (S26). Third, they do not require that the firing rates be positive, whereas we naturally enforce positivity through the transformation Eq. (S7).

#### A.9 Weighted versions of the problem

Under the assumption of a uniform prior, the formulation of our question treats all pairs of *θ*’s equally important. In natural settings or controlled experimental designs, this may not be the case. For example, horizontal and vertical orientations may occur more frequently in natural scenes [72], making *p*(*θ*) nonuniform. Then the pair (*θ, θ*^′^) to be distinguished occurs at frequency *p*(*θ*)*p*(*θ*^′^), and the cost function becomes

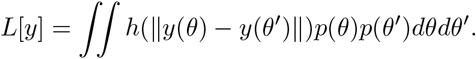

At the same time, previous works suggest that a signature of efficient coding (in the sense of maximizing mutual information) is the square root of 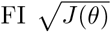 being proportional to the prior [21, 29]. Then the constraint on the curve *y*(*θ*) becomes

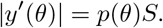

We can rescale the parameterization of the curve to recover the original constant-speed constraint. Consider the arclength parameterization

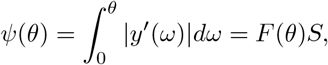

where *F* (*θ*) is the cdf of *p*(*θ*). Then using the inverse function theorem we have

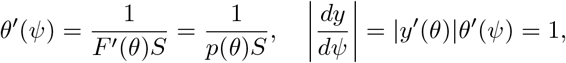

which states that *y*(*ψ*) has constant speed. The cost function becomes

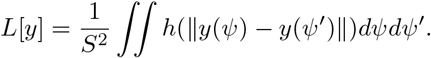

That is, we recover the original problem Eq. (S9) under the arclength parameterization. Then the optimizer is *y*(*ψ*) being a unit-speed circle, and the curve *y*(*θ*) in the original stimulus parameterization becomes a redistribution of points on the circle according to the efficient coding principle FI ∝ *p*(*θ*) [21, 29].

More generally, we can consider the joint distribution of the pair (*θ, θ*^′^) beyond the marginal:

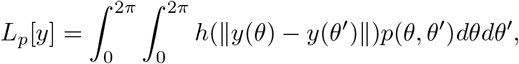

where *p*(*θ, θ*^′^) is the probability of the pair (*θ, θ*^′^) appearing in a task that requires downstream neurons to distinguish between them. One natural assumption on *p*(*θ, θ*^′^) is that it depends on the distance *d*(*θ, θ*^′^) (see Eq. (S27) for the appropriate definition of distance on *S*^1^) and remains uniform in the marginals *p*(*θ*) and *p*(*θ*^′^). For example, discriminating between *θ* and *θ*^′^ that are too close together may no longer provide behavioral benefits, and *p*(*θ, θ*^′^) is close to 0 for small *d*(*θ, θ*^′^). Notice that this distance and the distance *d*_*y*_ along the curve *y*(*θ*) only differ by a scaling constant. Then the full form of theorem 1 that includes a dependence of *g* on *d*_*y*_ can be used to prove that the solution is again the round circle.

#### A.10 Extension to hyperspheres *M* = *S*^***m***^ **and hypercubic flat hypertori *M* = *T*** ^***m***^

In this section we consider extensions of theorem 1 to the case where the stimulus *θ* lies on an *m*-sphere of radius 1 (i.e., *M* = *S*^*m*^) with *m < N* or a hypercubic flat *m*-torus (i.e., *M* = *T* ^*m*^ = ℝ^*m*^*/*(2*π*ℤ)^*m*^) with 2*m < N* . We only prove the simple versions (without the second term of *g*) for brevity, but the general versions are easily obtainable.

In the following we will use the assumption that an embedding *y* : *M* → ℝ^*N*^ is a short map. This means

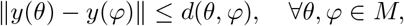

where *d*(·, ·) is the geodesic distance on the manifold *M* . Note that it in particular implies ∥*dy*_*θ*_∥_2_ ≤ 1, ∀*θ* ∈ *M*, where *dy*_*θ*_ is the differential of *y* at *θ*.

##### Theorem 3.

*Let y* : *S*^*m*^ → ℝ^*N*^ *be an embedding that is a short map. For a real function g*(*d*) *that is strictly convex and strictly decreasing in d*^2^, *the energy*

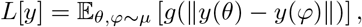

*with θ, φ independently drawn from the uniform distribution µ on S*^*m*^, *is uniquely minimized by inclusion in the sense that y is isometric with y*(*S*^*m*^) *a m-sphere contained in a* (*m* + 1)*-dimensional affine subspace of* ℝ^*N*^.

*Proof*. Let us first prove a Wirtinger-type inequality as in [46]. Define

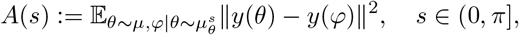

where 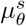 is the uniform distribution of points *φ* ∈ *S*^*m*^ that satisfy *d*(*θ, φ*) = *s*. This set is an intersection of two *m*-spheres and is itself a (*m* − 1)-sphere, so a uniform distribution on it is well defined. That is, *A*(*s*) is the average squared Euclidean distance between random pairs of embedded points coming from points on *S*^*m*^ with geodesic distance *s*. We can decompose the expectation into two steps: first take a random great circle *C* ⊂ *S*^*m*^, then scan over all pairs of points on the circle. Let *c*(*t*) be the arclength parameterization of a sample from *C*. Due to the symmetry of *S*^*m*^, taking a random great circle can be easily formulated as taking a random two-dimensional subspace from the Grassmannian. This recovers the one-dimensional circle case already presented in theorem 5 of [46]. We have

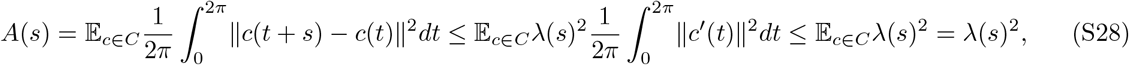

where *λ*(*s*) := 2 sin(*s/*2). Here the first inequality is due to theorem 5 of [46]; the second inequality is due to ∥*c*^′^(*t*)∥ ≤ 1 which is a consequence of *y* being a short map; and the last equality is due to the bound *λ*(*s*) being independent of the great circle *c* ∈ *C*. We can then proceed to use Jensen’s inequality with *g*’s strict monotonicity and convexity w.r.t. the square of its argument and to get

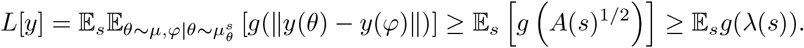

The lower bound is clearly attained by an inclusion. In the other direction, attaining the lower bound implies that Jensen’s inequality becomes an equality for all *s*, which requires ∥*y*(*θ*) − *y*(*φ*) ∥ ≡ *λ*(*d*(*θ, φ*)) for all pairs *θ, φ*. Specifying the pairwise distance of all embedded points uniquely determines the embedding up to rigid motion, so the map has to be an inclusion.

Note that the round sphere has been proven to uniquely minimize the global tangent-point energy, which is another form of nonlocal cost function [114].

##### Theorem 4.

*Let y* : *T* ^*m*^ → ℝ^*N*^ *be an embedding that is a short map. Here T* ^*m*^ = ℝ^*m*^*/*(2*π*ℤ)^*m*^ *is the flat hypertorus obtained by gluing opposite faces together. For a real function g*(*d*) *that is strictly convex and strictly decreasing in d*^2^, *the energy*

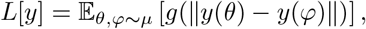

*with θ, φ independently drawn from the uniform distribution µ on T* ^*m*^ *(defined using the Lebesgue measure), is uniquely minimized by inclusion in the sense that y is isometric with y*(*T* ^*m*^) *the product of m round circles*.

*Proof*. We again want to prove a Wirtinger-type inequality. However, we can no longer invoke theorem 5 of [46] directly, as geodesics on *T* ^*m*^ are not identical circles as the spherical case. Fortunately, *T* ^*m*^ is naturally amenable to Fourier decomposition, and we only need to modify the Fourier series proof of [46]’s theorem 5 to handle “oblique” segments. The key is to utilize the reflection symmetries of *T* ^*m*^ and group Fourier terms by sign.

Let the Fourier series of *y*(*θ*) be

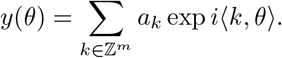

Then using the orthonormality of the Fourier basis we have

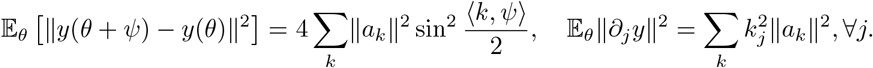

The Wirtinger-type inequality in theorem 5 of [46] is stated for each segment length on the circle. Here we cannot relate the above two quantities for each displacement *ψ*, but this becomes possible when we average the signs of *ψ*’s entries, which is sufficient for our purposes since we will eventually average over *ψ* to prove the final result. Concretely, let *ξ* ∈ {−1, 1}^*m*^ and define *ψ*^*ξ*^ := (*ξ*_1_*ψ*_1_, …, *ξ*_*m*_*ψ*_*m*_). The coordinate reflection symmetries of *T* ^*m*^ suggest that we only need to prove a Wirtinger-type inequality for averages over *ξ*. Define

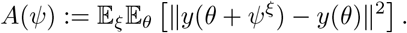

Recall the general trigonometric identity

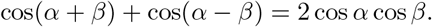

This allows us to calculate

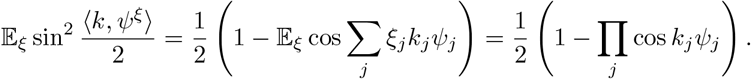

Recall that for |*γ*_*j*_| ≤ 1, ∀*j*, we have the general inequality

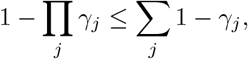

with equality if and only if at most one *γ*_*j*_ *<* 1. Then

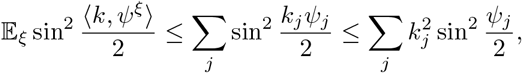

where we used lemma 6 of [46] in the second inequality. For generic *ψ*_*j*_’s (i.e., those that are not multiples of *π*), we have 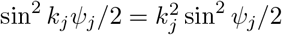 if and only if *k*_*j*_ = 0, ±1. We then have

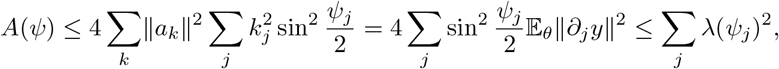

where *λ*(*s*) := 2 sin(*s/*2) and we used the assumption that *y* is a short map in the last inequality. Note that this is essentially a Pythagorean generalization of theorem 5 of [46], and equality is attained only by an inclusion. We can then apply Jensen’s inequality as we did in theorem 3 to conclude our proof.

The two proofs above can in principle be adapted and combined to generalize to direct products of rank-one symmetric spaces (with the hypertorus being a product of circles), but it is unclear how to generalize it further, as the decomposition of the integral in Eq. (S28) into circles of the same length is not possible for general manifolds.

Efficient coding requires 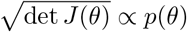 in higher dimensions [43, 44]. Here *J*(*θ*) becomes the Riemannian metric on the embedded manifold in *y* coordinates, and 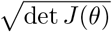 is the volume form. The arguments in Section A.9 can be directly adapted to the hypersphere and hypertorus cases using Moser’s trick [115].

The short map assumption in the above theorems is in particular satisfied by requiring *y* to be an isometric embedding, which can be interpreted as preserving the local discriminability relationship between stimuli. It is unclear whether neural representation necessarily needs to adhere to this constraint, but *some* constraint is needed in addition to pointwise volume (from efficient coding) to ensure that the optimization problem is well-posed when the stimulus is multi-dimensional. Minimizing the average binary classification error without any additional constraint generally leads to singular behavior such as stretching the sphere along an axis while thinning it along other axes to approach an infimum of the energy that is never attained.

We remark that the intrinsic (Riemannian) geometry of the stimulus manifold also becomes much richer when the stimulus has more than 1 dimension. For instance, there are different geometric classes of flat 2D tori, even though they are all topologically equivalent and locally flat. In 2D, our theorem 4 applies to square tori but not hexagonal tori, which more closely match the response pattern of grid cells. Considerations external to our framework are needed to differentiate square and hexagonal tori before asking what the optimal neural representation is for their respective intrinsic geometries.

### B Supplementary figures

**Fig. S1.**
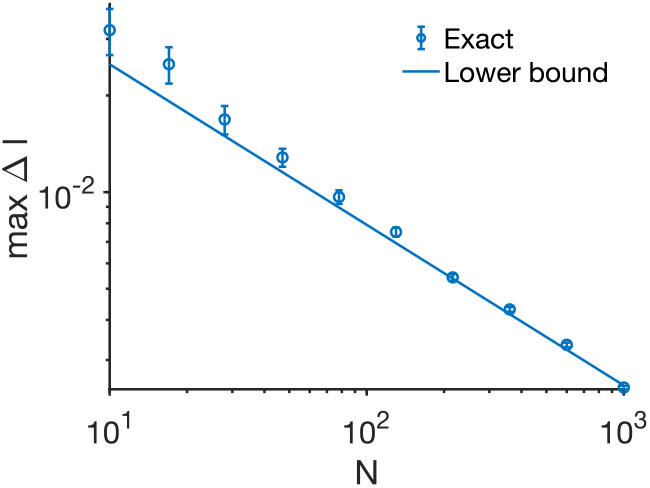
The lower bound given in Eq. (S14) compared with exact computation using Eq. (S12). Error bars are SEM over random circles *y*(*θ*).

**Fig. S2.**
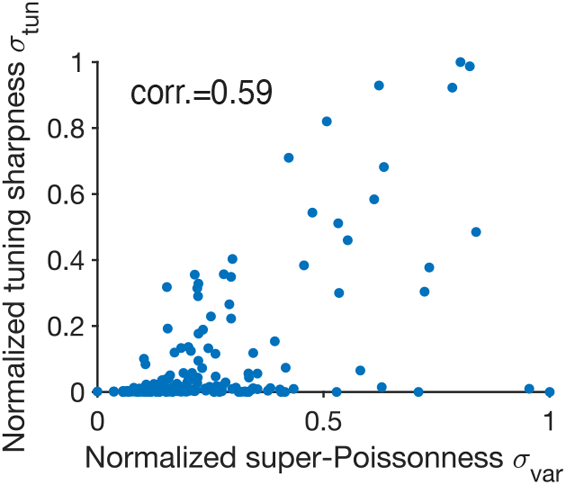
Same as Fig. 5c, but using data from monkey V1 during passive viewing of drifting gratings [30]. *N* = 169 neurons in total, combined across all 3 sessions.

